# Homeobox transcription factor MNX1 is crucial for restraining the expression of pan-neuronal genes in motor neurons

**DOI:** 10.1101/2021.08.07.455331

**Authors:** Ming-an Sun, Sherry Ralls, Warren Wu, Justin Demmerle, Jiayao Jiang, Carson Miller, Gernot Wolf, Todd S. Macfarlan

## Abstract

Motor neurons (MNs) control muscle movement and are essential for breathing, walking and fine motor skills. Motor Neuron and Pancreas Homeobox 1 (MNX1) has long been recognized as a key marker of the MN lineage. Deficiency of the *Mnx1* gene in mice results in early postnatal lethality – likely by causing abnormal MN development and respiratory malfunction. However, the genome-wide targets and exact regulatory function of *Mnx1* in MNs remains unresolved. Using an *in vitro* model for efficient MN induction from mouse embryonic stem cells, we identified about six thousand MNX1-bound loci, of which half are conserved enhancers co-bound by the core MN-inducing factors ISL1 and LHX3, while the other half are promoters for housekeeping-like genes. Despite its widespread binding, disruption of *Mnx1* affects the activity of only a few dozen MNX1-bound loci, and causes mis-regulation of about one hundred genes, the majority of which are up-regulated pan-neuronal genes with relatively higher expression in the brain compared to MNs. Integration of genome-wide binding, transcriptomic and epigenomic data in the wild-type and *Mnx1*-disrupted MNs predicts that *Pbx3* and *Pou6f2* are two putative direct targets of MNX1, and both are homeobox transcription factors highly expressed in the central nervous system. Our results suggest that MNX1 is crucial for restraining the expression of many pan-neuronal genes in MNs, likely in an indirect fashion. Further, the rarity of direct targets in contrast to the widespread binding of MNX1 reflects a distinctive mode of transcriptional regulation by homeobox transcriptional factors.

## Introduction

The spinal cord coordinates all muscle movement and consists of two types of neurons: motor neurons (MNs) that directly innervate and control muscle, and interneurons (INs) that mediate both descending signals from the brain and signal transduction between MNs and INs in the spinal cord (Jessell, 2000). Spinal cord MNs are essential for biological processes including breathing and movement, and dysfunction in MNs is associated with several human neuronal diseases such as amyotrophic lateral sclerosis and spinal muscular atrophy (Kiernan, 2018).

Spinal cord neurons are differentiated through a series of processes. First, different neural progenitor domains are formed in response to a dorsoventral Shh gradient produced by the notochord and floorplate of the neural tube, with each domain expressing distinct cross-repressive transcription factors that define domain boundaries. Neuronal progenitor cells (NPCs) differentiate into specific types of mature neurons under the control of a few core transcription factors (Jessell, 2000; Lee et al., 2008; Mazzoni et al., 2013). For example, the two LIM-homeobox transcription factors ISL1 and LHX3 together with their cofactor NLI (LDB1) form a 2NLI:2ISL1:2LHX3 hexamer complex that drives the specification of MNs (Thaler et al., 2002). NPCs in the pMN domain, a restricted domain of the ventral ventricular zone, differentiate into mature MNs after the MN-hexamer activates downstream genes (Lee et al., 2008; Thaler et al., 2002). Interestingly, the specification of one neuronal cell type is usually accompanied by the repression of the molecular characteristics of alternative cell types. For example, the expression of core transcription factors (eg. *Isl1*, *Lhx3* and *Mnx1*) during MN specification can repress V2a-IN genes such as *Vsx2* (Lee et al., 2012; Thaler et al., 1999). Reciprocally, the specification of V2a-INs is accompanied by the repression of MN genes such as *Mnx1* and *Isl1* (Clovis et al., 2016; Debrulle et al., 2019).

Motor Neuron and Pancreas Homeobox 1 (MNX1), also known as HLXB9 or HB9, is a conserved homeobox transcription factor expressed specifically during spinal MN specification, making it a widely used MN marker. Deficiency of *Mnx1* in mice causes early postnatal lethality, likely due to malfunction of the respiratory system (Arber et al., 1999; Thaler et al., 1999). Importantly, *Mnx1*-deficient mouse embryos still develop normal amounts of MNs, but these MNs develop inappropriate axonal projection, indicating *Mnx1* is essential for proper axon guidance (Arber et al., 1999; Thaler et al., 1999). The study of the *Mnx1* ortholog in Drosophila suggests that it controls axon guidance through indirect regulation of *Robo* receptors (Santiago et al., 2014). *Mnx1* has a conserved C-terminal homeobox domain that binds to DNA, and a N-terminal Engrailed Homology 1 (EH1) domain that can interact with the Groucho co-repressor (Copley, 2005). Luciferase reporter assays demonstrated that *Mnx1* and its chicken homolog *Mnr2* act as transcriptional repressors (Lee et al., 2008; William et al., 2003). However, until now only a few genes have been discovered to be mis-regulated after the disruption of *Mnx1*. The genome-wide targets and precise regulatory functions of MNX1 during MN specification remain to be systematically investigated.

MNs in the mouse spinal cord are of limited numbers and difficult to purify (Bjugn and Gundersen, 1993), and early *in vitro* MN induction methods from embryonic stem cells (ESCs) using the patterning factors Shh and retinoic acid (RA) are of relatively low efficiency (Wichterle et al., 2002). Recently developed techniques based on directed differentiation of MNs from ESCs by induced expression of *Ngn2, Isl1,* and *Lhx3* can produce large amounts of MNs with high purity (Mazzoni et al., 2013). Taking advantage of a similar model, in combination with endogenous epitope tagging and targeted mutagenesis of the *Mnx1* gene, we applied state-of-the-art OMICs techniques to investigate the molecular function of MNX1 during MN specification.

## Results

### High-efficient induction of motor neurons from mouse ESCs

We developed an *in vitro* model for directed induction of MNs from mouse ESCs (**Figure 1A**). Based on the previously developed A2lox mouse ESC line with inducible cassette exchange (Iacovino et al., 2011), a transgenic ESC line was developed with the following genetic modifications: 1) Insertion of NIL (*Ngn2+Isl1+Lhx3*) coding sequences in the *Hprt* locus, which enables doxycycline (DOX)-inducible overexpression of NIL to drive MN induction; 2) Insertion of a N-terminal eGFP coding sequence in-frame with MNX1, which facilitates fluorescence imaging and flow cytometry based on GFP signal to determine induction efficiency, and enables immunoprecipitation experiments using antibody against the GFP tag.

**Figure 1.**
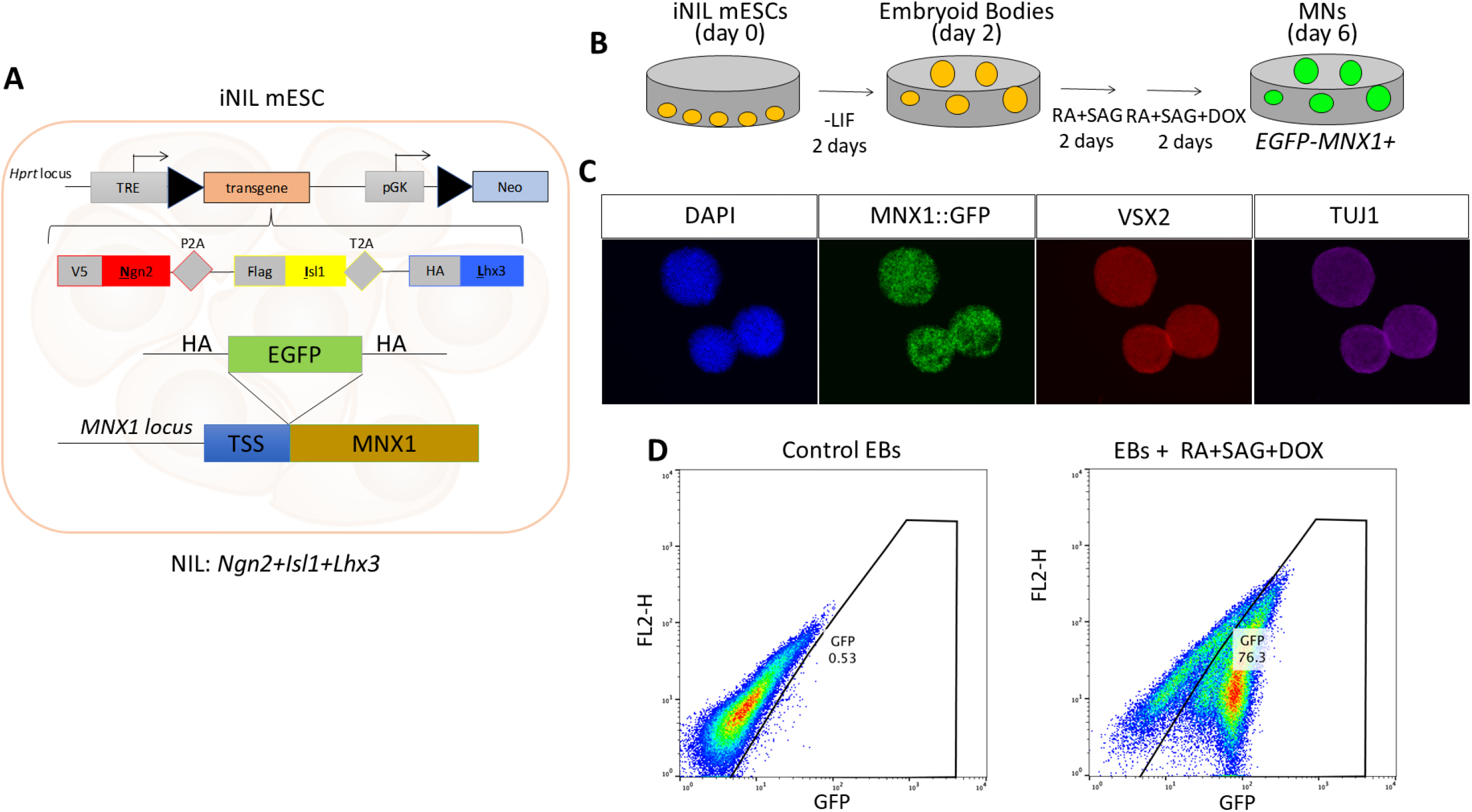
Directed motor neuron induction from mouse ESCs with high efficiency. **A.** Scheme for the transgenic mESCs used in this study, with features including: 1) DOX-inducible NIL coding sequence inserted downstream of the *Hprt* locus; 2) eGFP coding sequence inserted immediately downstream of the *Mnx1* promoter. **B.** The 6-day procedure for directed MN induction from mESCs. **C.** Immunofluorescence imaging for DAPI, GFP, VSX2 and TUJ1 in induced MNs. The expression of *Mnx1* is demonstrated by the MNX1::GFP signal. **D.** FACS results demonstrating iMNs express MNX1::GFP with high efficiency.

The MN induction was performed following a 6-day procedure (**Figure 1B**) similar to previous studies (Mazzoni et al., 2013; Mazzoni et al., 2011). In brief, mouse ESCs are cultured for two days as embryoid bodies on a shaking platform in neural differentiation media, followed by two days of differentiation in the presence of RA and Smoothened agonist (SAG). On the fourth day, the media is replaced with fresh RA and SAG along with DOX to induce the expression of the NIL transgene. Fluorescence imaging and FACS demonstrate that the induction efficiency is typically 75% by day 6 but varies between 60%-90% (**Figure 1C,D**). To minimize batch effects, we limited downstream experiments to induced MNs (iMNs) produced at efficiencies greater than 70%. The RNA-Seq and ChIP-Seq data generated from the mouse ESCs and iMNs and collected from previous studies are summarized in **Table S1**.

### MNX1 binds thousands of conserved neuronal enhancers and housekeeping-like promoters

To determine the direct targets of MNX1, we profiled its genome-wide binding in wild-type iMNs using ChIP-Seq with an antibody recognizing the in-frame GFP tag. Our data confirm MNX1 binding near two known target genes, *Mnx1* itself and *Vsx2*, indicating the good quality of our data (**Figure S1**). In total, we identified 5,943 MNX1-bound loci, with 40.8% (n = 2,419) located within promoters, which is remarkably higher than expected by random (**Figure 2A**). Of the remaining loci, 26.0% and 29.2% are within intronic and intergenic regions, respectively. MNX1-bound loci are highly conserved, and interestingly those in intronic or intergenic regions have even higher sequence conservation than those in promoters (**Figure 2B**). Given that sequence conservation can be used to predict regulatory elements (King et al., 2005), we speculate that the bound intronic and intergenic loci are putative enhancers. We also profiled the occupancies of five types of histone marks in iMNs, which confirm that MNX1-bound intronic and intergenic loci are enriched with H3K27ac and H3K4me1, which are known to mark active and/or poised enhancers (**Figure 2C**). The MNX1-bound loci in promoter regions are highly enriched with H3K27ac and H3K4me3, which usually occur on active promoters. Despite *Mnx1* being reported as a transcriptional repressor (Lee et al., 2008; William et al., 2003), its bound loci are more frequently associated with active histone marks (H3K4me3, H3K27ac and H3K4me1) and not repressive marks like H3K27me3 and H3K9me3 (**Figure 2C, S1**).

**Figure 2.**
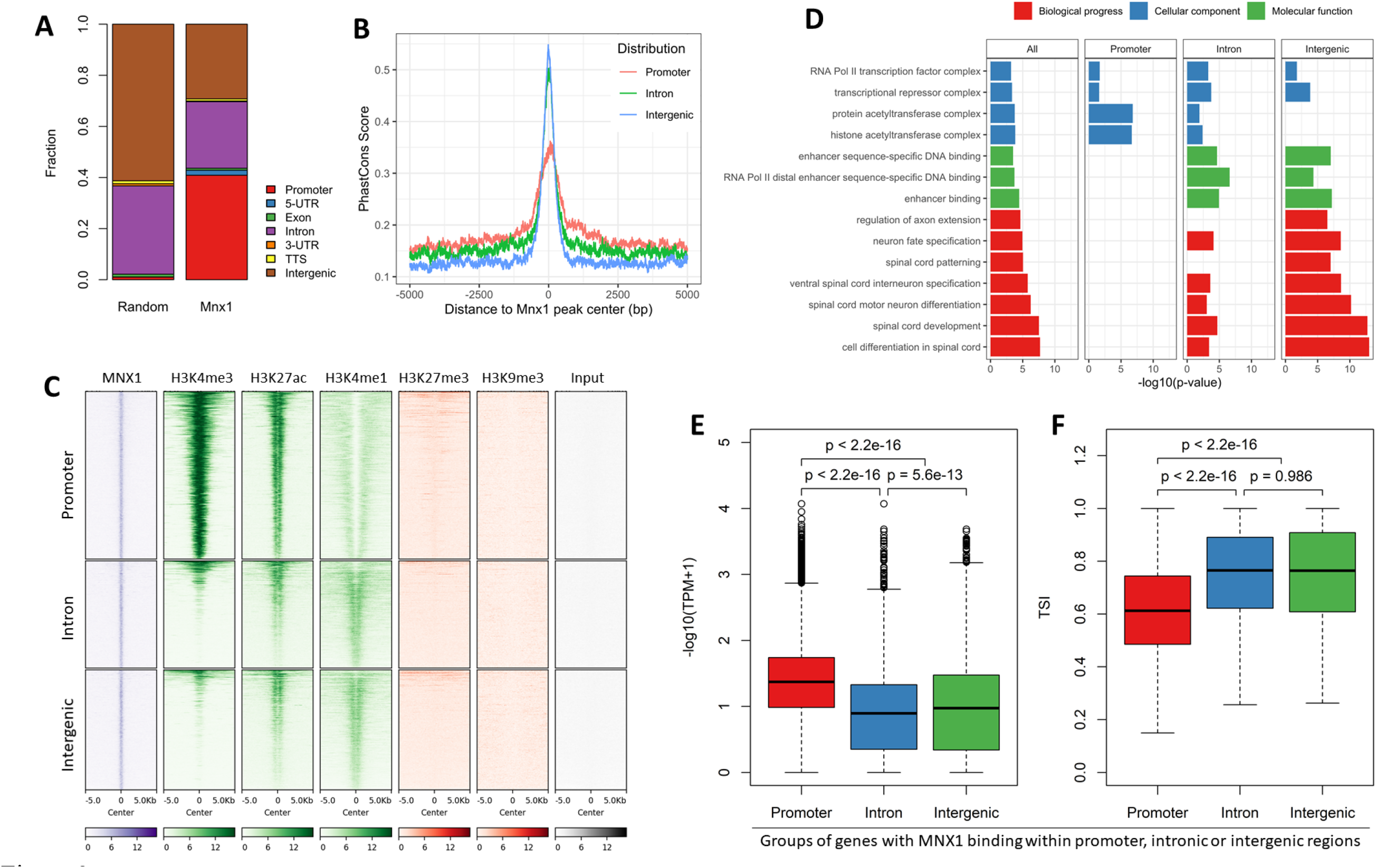
Binding preference of MNX1 on thousands of conserved enhancers and promoters. **A.** Genomic distribution of MNX1-bound loci relative to randomly selected genomic regions. **B.** Conservation of MNX1-bound loci in promoters, intronic or intergenic regions. **C**. Heatmaps for the occupancy of different histone marks surrounding MNX1-bound loci within promoters, intronic or intergenic regions. **D.** Enriched GO terms associated with different groups of MNX1-bound loci based on their genomic distribution. The top 1000 loci are used for analysis. Representative terms from three GO categories are visualized in the barplots. **E,F**. Boxplots showing genes within MNX1 binding in their promoters are usually of high expression level (**E**) and low tissue specificity (**F**).

We performed Gene Ontology (GO) enrichment analysis on MNX1-bound loci and found they are indeed highly associated with neuronal functions, particularly spinal cord development/patterning, motor neuron and interneuron specification (**Figure 2D**). However, closer inspection showed that only MNX1-bound loci within intronic and intergenic regions are associated with neuronal functions, while those in promoters are associated with more general functions such as non-coding RNA metabolic process, RNA transport and protein folding (**Figure 2D**). Further examination of the transcriptomic data in iMNs and twelve other mouse tissues showed that genes associated with MNX1-bound promoters are of relatively high average expression level and low tissue specificity, a signature of housekeeping genes (**Figure 2E**). In contrast, genes with MNX1 bound in their introns or flanking intergenic regions have relatively low expression levels and high tissue specificity (**Figure 2F**). Taken together, our results suggest that MNX1 binds both promoters and enhancers, yet only the enhancers are associated with neuronal genes, while the promoters tend to be associated with housekeeping-like genes.

### MNX1 colocalizes with the core MN inducing factors ISL1/LHX3 on thousands of neuronal enhancers

The NIL factors (Ngn2, Isl1 and Lhx3) play vital roles for MN specification, and their forced expression can directly induce MNs from mouse ESCs (Mazzoni et al., 2013). Previous profiling of NIL binding during MN induction demonstrated that ISL1 and LHX3 have almost identical binding profiles on motor neuron enhancers (Velasco et al., 2017). To determine how the binding of MNX1 and ISL1/LHX3 are related in iMNs, we retrieved the ChIP-Seq data of LHX3 from a previous study (Velasco et al., 2017) and compared it to the genomic binding sites of MNX1. K-mean clustering (k=2) of MNX1-bound loci based on the occupancy of MNX1, LHX3, and five histone marks confirmed the binding of MNX1 at putative both promoters (Cluster 1) and enhancers (Cluster 2), as indicated by their respective histone marks and genomic distributions (**Figure S2A-C**). However, the majority (96%) of LHX3-bound loci are in introns or intergenic regions but not in promoter regions (**Figure S2D**). Further comparison of MNX1 and LHX3 binding demonstrated that about half (48%) of MNX1-bound enhancers are co-occupied by LHX3 (**Figure S2E,F**).

To further characterize the putative regulatory elements with MNX1 and/or LHX3 binding, we classified them into four groups: group 1 consisted of MNX1-bound promoters, while groups 2, 3 and 4 were designated as enhancers bound by MNX1 alone, MNX1+LHX3, or LHX3 alone, respectively (**Figure 3A,B**). All three groups of enhancers, but particularly those co-bound by MNX1+LHX3, are highly conserved and associated with strong H3K27ac and H3K4me1 occupancy (**Figure 3A–C**). Motif analyses indicate that all three groups of enhancers are enriched with AT-rich homeobox binding motifs, but each with distinctive preference: enhancers bound with MNX1+LHX3 (G3) or LHX3 (G4) are enriched with the known LHX3 motif, while enhancers bound by MNX1 alone (G2) are enriched with motifs for BARX1, PDX1 and NKX3-1, which belong to homeobox genes or their co-factors (**Figure 3D, Table S2**). These enriched motifs also match the known MNX1 motifs identified by high-throughput SELEX (**Figure 3E**) in previous studies (Jolma et al., 2013; Zhu et al., 2018). In contrast, MNX1-bound promoters are enriched with motifs for SP2, NFYA and NRF instead of known MNX1 motifs (**Figure 3D**). GO enrichment analyses demonstrate that all groups of enhancers are associated with neuronal functions such as neuron differentiation and axon guidance, yet loci co-bound by MNX1+LHX3 usually have more significant p-values (**Figure 3F**). Together, our results suggest that MNX1 binds thousands of enhancers, and that about half are enhancers co-occupied by the core neuronal factors LHX3 and ISL1.

**Figure 3.**
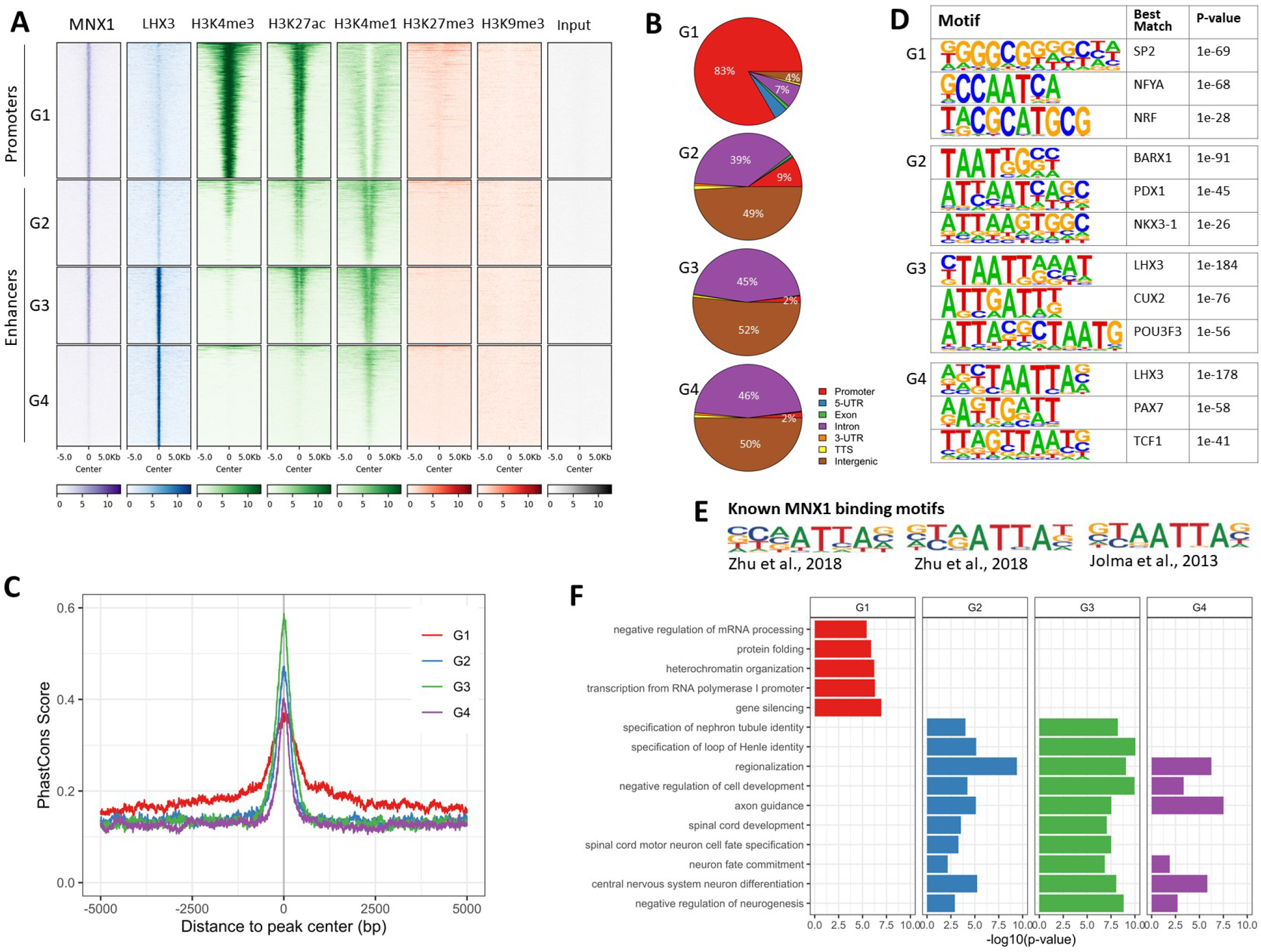
MNX1 and LHX3 co-bind thousands of conserved neuronal enhancers. **A.** Heatmaps for the occupancy of MNX1, LHX3 and different histone marks on four groups of loci bound by MNX1 and/or LHX3 (labelled as G1, G2, G3, or G4). Group 1 is MNX1-bound promoters, and groups 2-4 are enhancers bound by MNX1, MNX1+LHX3, or LHX3, respectively. These groups remain the same throughout this figure. **B.** Genomic distributions of different groups of loci bound by MNX1 and/or LHX3. **C**. Sequence conservation as measured by Phastcons score flanking different groups of loci bound by MNX1 and/or LHX3. **D**. Representative motifs enriched in loci bound by MNX1 and/or LHX3. **E.** Known MNX1 binding motifs reported in previous studies**. F**. GO enrichment for different groups of loci bound by MNX1 and/or LHX3.

### Disruption of *Mnx1* causes mis-regulation of dozens of pan-neuronal genes

Previous studies in mice suggest that disruption of *Mnx1* affects axon guidance, yet until now few genes, including *Vsx2* and *Isl1*, have been validated as mis-expressed in *Mnx1* mutants (Arber et al., 1999; Thaler et al., 1999). *Mnx1* has two conserved functional domains: a N-terminal homeodomain (HD) that binds DNA, and a C-terminal EH1 domain (EH1) that interacts with Groucho-like co-repressors (**Figure S3**). Using CRISPR/Cas9-based gene editing, we generated mutant mESC lines with either HD or EH1 deletions (**Figure 4A**). We generated four mutant lines in total, including one with an in-frame HD deletion (*Mnx1*ΔHD), one with a HD deletion causing a frameshift that results in early translational termination and complete degradation of MNX1 protein (*Mnx1*-null), and two replicate lines (*Mnx1*ΔEH1-1 and *Mnx1*ΔEH1-2) with in-frame EH1 deletions (**Figure S4, S5**). By culturing the iMNs collected at day 6 for two more days following an axon elongation protocol (Wu et al., 2012), we observed normal axon elongation in both *Mnx1*-null and *Mnx1*ΔHD mutants (**Figure S6**).

**Figure 4.**
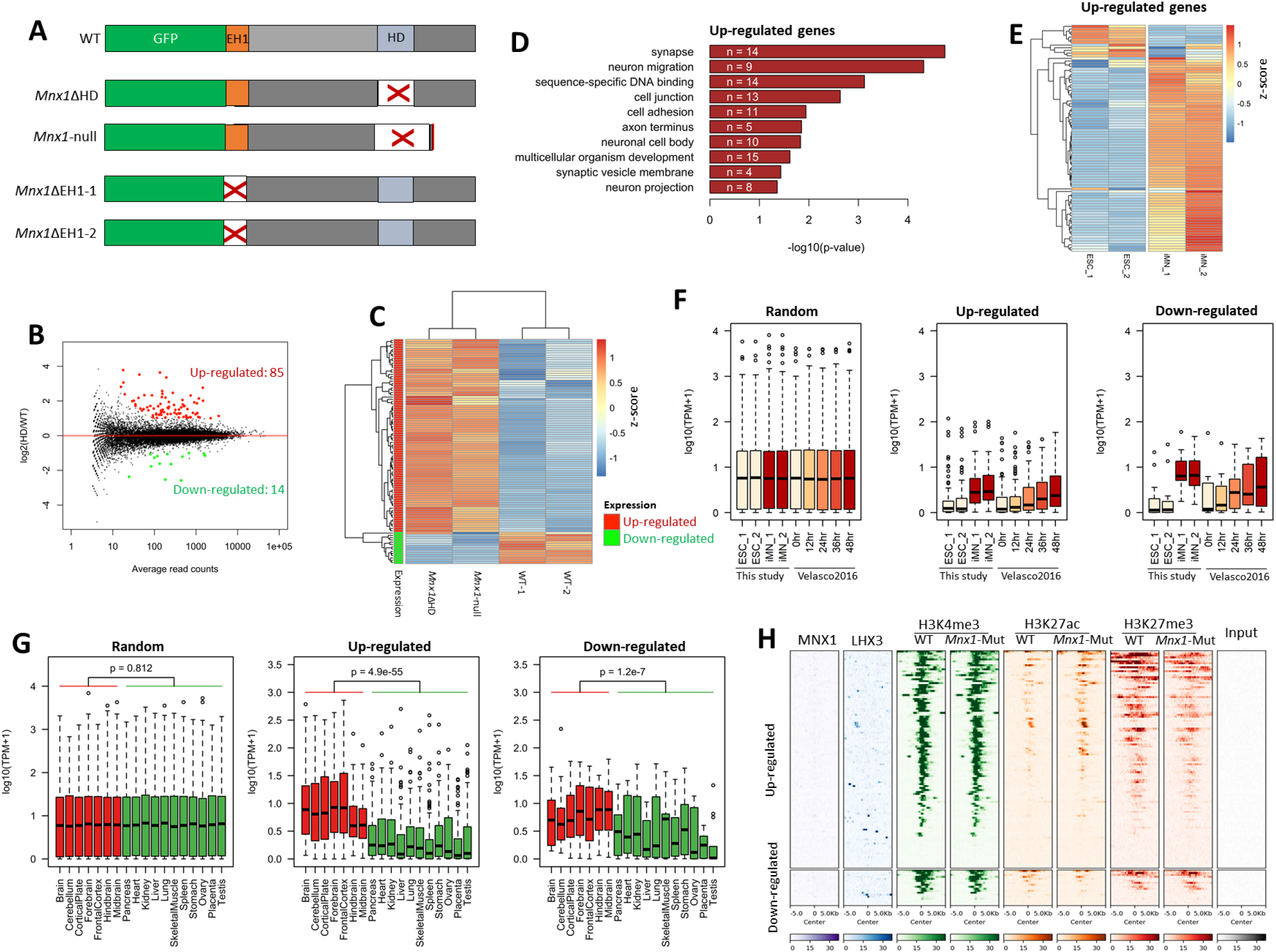
Disruption of *Mnx1* causes mis-regulation of dozens of neuronal genes. **A.** Scheme for the deletion of Homeodomain (HD) and EH1 motifs (EH1) from *Mnx1*. The EH1 motif and homeodomain are represented as orange and light blue boxes, respectively. The deleted region in each mutant is indicated as a white box with a red cross inside. Of note, one mutant with HD deletion (*Mnx1-*null) has a frameshift which then creates new stop codon for early termination. The stop codon created is indicated as a red bar. This mutant line is denoted as *Mnx1*-null due to the degradation of the MNX1 protein. **B.** MA-plot for the differential gene expression between *Mnx*1 mutants (*Mnx1*ΔHD and *Mnx1*-null) and WT iMNs. **C**. Expression profiles of the identified DEGs in different *Mnx1* mutants and WT iMNs. **D**. Enriched GO terms associated with the up- regulated genes in *Mnx1* mutants. The numbers of genes associated with each term are indicated in the plot. **E**. Expression profiles of the up-regulated genes in WT ESCs and iMNs. **F**. Expression profile of the up- and down-regulated genes during MN induction. Apart from the DEGs, 1000 randomly selected genes are used as a control. **G**. Expression profiles of the up- and down-regulated genes among different tissues. The expression data are retrieved from the ENCODE project. P-values are calculated using a two-tailed Student’s t-test. **H**. Occupancy of MNX1, LHX3 and different histone marks in WT and *Mnx1* mutants flanking the promoters of DEGs.

To understand the function of *Mnx1* on global gene expression, we generated RNA-Seq data for wild-type and *Mnx1* mutant iMNs. The RNA abundance of *Mnx1* remains similar between wild-type and different *Mnx1* mutants (**Figure S7**). We compared the *Mnx1*-null with *Mnx1*ΔHD and *Mnx1*ΔEH1 mutant iMNs to assess the contribution of each domain in regulating gene expression, and found that the *Mnx1*ΔHD and *Mnx1*-null mutations have similar effects on global gene expression, indicating that the homeodomain is essential for *Mnx1* function (**Figure 4C, S8, S9A**). In contrast, *Mnx1*ΔEH1 mutant iMNs show significant yet weakly correlated effects compared with the *Mnx1*-null mutant, possibly because EH1-deletion cannot fully disrupt the function of *Mnx1* (**Figure S9B**). We identified 99 differentially expressed genes (DEGs) by comparing *Mnx1*ΔHD and *Mnx1*-null mutants to *Mnx1* wild-type, with the majority (n=85) up-regulated (n=85) (**Figure 4B,C**). The up-regulated DEGs are highly associated with neuronal functions including synapse, neuron migration and axon terminus (**Figure 4D**). Indeed, many up-regulated genes are well-known neuronal genes, such as *Reln*, *Onecut1*, *Lmo3*, *Robo1*, *Ntng1*, *Pbx3*, *Synp*r, *Pcdh17*, *Syt4/6*, *Sorcs3*, *Foxp1* and *Phox2b* (**Figure S10**). While *Vsx2,* which is known to be repressed by MNX1, does not show up in the DEG list, close inspection confirms that *Vsx2* expression is increased by more than two-fold (log2fold=1.25, adjusted-p-value=0.36). Interestingly, most DEGs – irrespective of the direction of change – are activated during MN induction (**Figure 4E,F, S11**). Not surprisingly, examination of gene expression in iMNs and 18 mouse tissues demonstrates that most DEGs are highly expressed in the central nervous system (**Figure 4G, S11**). Consistent with expression changes, these DEGs usually have altered H3K27ac and H3K27me3 in their promoter regions (**Figure 4H, S12**). Together, disruption of *Mnx1* causes differential expression of almost one hundred genes, with most of those being up-regulated during MN induction. However, those DEGs are usually not MN-specific, as many are pan-neuronal genes with similar or even higher expression in brain regions relative to iMNs.

### *Pbx3* and *Pou6f2* are identified as the putative direct targets of *Mnx1*

Despite widespread binding on almost six thousand loci, disruption of *Mnx1* only alters the expression of ninety-nine genes in iMNs – indicating only a small fraction of the bound loci are under tight regulation by MNX1. This is further supported by two observations. First, various histone modifications on these loci remain largely unaltered after disruption of *Mnx1* (**Figure S13)**. Second, the genes mis-expressed after disruption of *Mnx1* are not enriched for MNX1 binding in their promoters or flanking regions, indicating that they are indirectly affected by the loss of MNX1 (**Figure 4H**). Identifying the enhancers and genes under direct regulation of *Mnx1* is therefore key to understand its function. Given that MNX1 interacts with the Groucho co-repressor, which interacts with HDACs (Chen et al., 1999; Winkler et al., 2010), we designed an unbiased computational procedure to infer *Mnx1*-regulated enhancers and genes by integrating genome-wide binding, transcriptomic and epigenomic data in wide-type and *Mnx1*-mutant iMNs (**Figure 5A**). In brief, we first identify putative *Mnx1*-targeted cis-elements as those loci with differential H3K27ac occupancy between WT and *Mnx1* mutants (*i.e. Mnx1*ΔHD and *Mnx1*-null) iMNs. Then, DEGs between WT and *Mnx1*-mutant iMNs were identified and compared against the differential H3K27ac loci. Finally, the putative *Mnx1* target genes were identified as DEGs associated with at least one H3K27ac loci significantly changed at the same direction.

**Figure 5.**
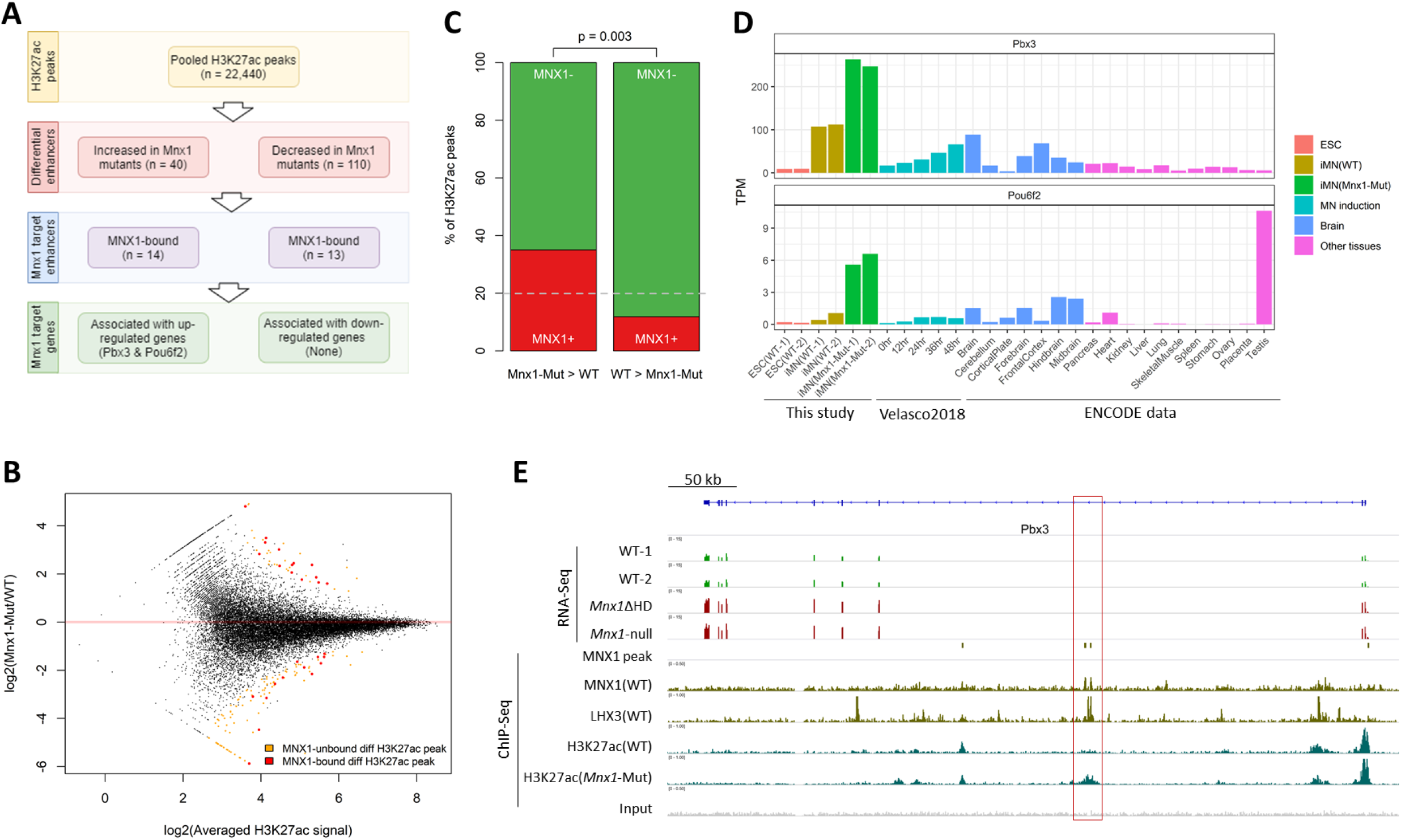
Identification of *Pbx3* and *Pou6f2* as putative direct targets of MNX1. **A.** Flowchart for the computational inference of putative direct targets of MNX1. **B.** Differential H3K27ac occupancy between WT and *Mnx1*-mutant iMNs. Significant differential peaks are highlighted in orange, and among them, those bound by MNX1 are highlighted in red. **C.** Fraction of MNX1-bound loci for significantly increased and decreased H3K27ac peaks, respectively. Loci bound or not bound by MNX1 are red and green, respectively. The grey dashed line shows the averaged fraction of all H3K27ac peaks bound by MNX1. P-values are calculated with Fisher’s Exact Test. **D**. Expression pattern of *Pbx3* and *Pou6f2* in different cell types (ESC and iMN), during MN induction (0 hr – 48 hr) and among different tissues. **E**. IGV tracks for the expression abundance and occupancy of MNX1, LHX3 and H3K27ac near the *Pbx3* locus for WT and *Mnx1*-mutant iMNs. The putative *Mnx1*-regulated locus is highlighted by the red rectangle.

Of the 22,440 pooled H3K27ac peaks identified in WT and *Mnx1*-mutant iMNs, only 150 (0.67%) were altered in *Mnx1*-mutants, including 40 loci with increased H3K27ac and 110 loci with decreased H3K27ac (**Figure 5A,B**). MNX1 binding is enriched at H3K27ac-increased loci and depleted at H3K27ac-decreased loci (odd-ratio = 2.96, P-value=0.003; **Figure 5C**), in agreement with previous studies suggesting that MNX1 is a transcriptional repressor (Lee et al., 2008; William et al., 2003). We found only 14 H3K27ac-increased loci and 13 H3K27ac-decreased loci that are bound by MNX1, and of the H3K27ac-decreased loci none are associated with changes in gene expression. In contrast, two H3K27ac-increased loci in *Mnx1* mutants reside in introns of two up-regulated genes, *Pbx3* and *Pou6f2,* both of which are co-bound by LHX3 (**Figure 5D, S14**). No other candidates were identified even after allowing the H3K27ac-increased loci to be as far as 50 kb away from up-regulated genes. Both *Pbx3* and *Pou6f2* are homeobox transcription factors that are gradually expressed during MN induction and are overexpressed in *Mnx1* mutants (**Figure 5D,E**). Further examination indicates that both *Pbx3* and *Pou6f2* are highly expressed in the central nervous system, particularly *Pbx3* which is highly expressed in iMNs relative to brain and other tissues (**Figure 5D**). Together, our results indicate that the direct genetic targets of *Mnx1* are rare, and that *Pbx3* and *Pou6f2* are the two most plausible direct targets even though their roles in MN development remain to be clarified.

## Discussion

Homeodomain proteins have indispensable roles in various developmental processes, including neuronal differentiation and specification (Burglin and Affolter, 2016; Reilly et al., 2020). Among them, *Mnx1* is specifically expressed in spinal MNs and has been widely used as a MN marker. Previous studies suggest that *Mnx1* and its orthologs regulate axon guidance, and disruption of *Mnx1* causes early postnatal lethality in mice (Arber et al., 1999; Santiago et al., 2014; Thaler et al., 1999). However, while it is recognized that MNX1 binds and represses the interneuron gene *Vsx2 (Arber et al., 1999; Thaler et al., 1999)*, its binding targets and regulatory mechanisms remain largely unknown. Taking advantage of an *in vitro* MN induction procedure together with CRISPR engineering and multi-dimensional OMICs profiling, we systematically investigated the regulatory elements and genes that are potentially under the regulation of *Mnx1*.

We determined the genome-wide binding of MNX1 in iMNs and found that it binds thousands of conserved promoters and enhancers. The bound enhancers are associated with neuronal functions, which is not surprising given the known importance of *Mnx1* in MNs (Arber et al., 1999; Thaler et al., 1999). Indeed, more than half of MNX1-bound enhancers are co-bound by the core MN-inducing transcription factors ISL1 and LHX3, implying their potential cooperation at these loci. But unlike ISL1 and LHX3 which exclusively bind enhancers (Velasco et al., 2017), MNX1 also binds thousands of promoters associated with house-keeping-like genes. Binding of MNX1 at these seemingly irrelevant promoters is unexpected, but the MNX1 ChIP-Seq data are reliable given that they are generated using an antibody that recognizes an endogenous GFP tag, and that binding near two known targets of MNX1, *Mnx1* and *Vsx2,* are confirmed. Given that only MNX1-bound enhancers, but not MNX1-bound promoters, show overrepresentation of the TAATTA motif resembling the one determined by a SELEX-based assay for MNX1 binding (Jolma et al., 2013; Zhu et al., 2018), we speculate that MNX1 binding in promoters is through a different mechanism relying on co-factors instead of its homeodomain. How and why MNX1 binds numerous house-keeping-like promoters in iMNs is yet to be determined, and it is also unclear if this occurs with other homeobox transcription factors.

MNX1 has long been considered as a transcriptional repressor, with its repressive function likely mediated by Groucho proteins (Arber et al., 1999; Lee et al., 2008; Thaler et al., 1999; William et al., 2003). However, disruption of *Mnx1* only alters the expression of about one hundred genes, and the activity of only a few dozens of MNX1-bound loci, which contrasts with its widespread binding on almost six thousand loci in total. There are several possible explanations. Firstly, is possible that only a small percentage of the MNX1-bound loci are under the tight regulation of *Mnx1* – at least for the *in vitro* MN induction model used in this study. This would be not surprising given that many homeobox transcription factors recognize similar AT-rich motifs (Berger et al., 2008; Jolma et al., 2013; Langmead and Salzberg, 2012; Zhu et al., 2018). It is possible that co-binding of MNX1 and other homeobox transcription factors (*e.g.* ISL1 and LHX3) may be functionally redundant on some loci. Secondly, our *in vitro* model studies nearly pure MN populations at relatively early developmental stages, which is not identical to *in vivo* MNs which consist of various cell subtypes that continue to express *Mnx1* after specification. Therefore, we cannot exclude the possibility that MNX1 regulates specific MN subtypes which cannot be monitored using an *in vitro* MN induction model. Lastly, it is also possible that the co-repressors necessary for MNX1 function are absent from many bound loci. For example, Groucho proteins are known to recruit histone deacetylases (HDACs) to mediate transcriptional repression (Arce et al., 2009; Sekiya and Zaret, 2007). Since most MNX1-bound loci have active histone modifications that remain unaltered after the disruption of *Mnx1*, it is possible that Groucho co-repressors are absent from most of these MNX1-bound loci. It would be informative to profile the binding of two Groucho orthologs that have increased expression during MN induction, TLE1 and TLE3, but this was failed with ChIP-Seq using commercial antibodies.

Disruption of *Mnx1* alters the expression of about one hundred genes, with the majority being up-regulated. While most of these DEGs are neuronal genes that are induced during MN differentiation, surprisingly most of them are not specifically expressed in MNs. The mis-regulated genes caused by disruption of *Mnx1* are usually pan-neuronal genes with similar or even higher expression in other parts of the central nervous system (*i.e.* different regions of the brain) relative to MNs. Why does *Mnx1* seem to regulate pan-neuronal genes which are not specific for MNs? We speculate that *Mnx1* behaves like a brake to restrain the expression of pan-neuronal genes in MNs to an appropriate level. Indeed, the previously reported transcriptional repression of the interneuron gene *Vsx2* by MNX1 (Lee et al., 2012; Thaler et al., 1999) could be considered as an extreme case following this hypothesis. By restraining the expression of pan-neuronal genes or genes for alternative neuronal types, *Mnx1* may reinforce the identity and function of MNs. However, unlike with *Vsx2* which is directly bound and regulated by MNX1, the effect of *Mnx1* on most of the mis-regulated pan-neuronal genes is likely to be indirect.

We designed an unbiased computational procedure to identify putative direct targets of *Mnx1* by integrating genome-wide binding, transcriptomic and epigenomic data in wild-type and *Mnx1*-mutant iMNs. Surprisingly, only two genes, *Pbx3* and *Pou6f2*, were predicted to be under the direct regulation of *Mnx1*. Interestingly, both *Pbx3* and *Pou6f2* are transcription factors that are highly expressed and have important roles in the central nervous system. *Pou6f2* is known to be expressed in distinct clades of V1 interneurons (Bikoff et al., 2016; Sweeney et al., 2018), where it has been predicted to be important for the trans-differentiation of excitatory projection neurons and inhibitory interneuron subtypes (Ainsworth et al., 2018). In addition, the *Pbx* genes have been recognized as key co-factors for *Hox* genes (Mann et al., 2009), and important for neuron differentiation (Grebbin et al., 2016; Schulte and Frank, 2014; Selleri et al., 2019). *Pbx3* and its paralogs *Pbx1/2/4* belong to the TALE homeobox family of transcription factors (Cerda-Esteban and Spagnoli, 2014), and among them *Pbx1* and *Pbx3* are highly expressed in mouse spinal MNs with distinct spatial distributions (Hanley et al., 2016). Yet, unlike *Pbx3* which is up-regulated after disruption of *Mnx1*, *Pbx1* is remarkably decreased, suggesting that *Pbx1* and *Pbx3* are under different modes of regulation (**Figure S15**). Previous studies suggest that *Pbx3*-deficient mice develop to term but die soon after birth from central respiratory failure (Onimaru et al., 2004; Rhee et al., 2004), and in the absence of *Pbx1*/*3* spinal MNs can still be formed, but the MN subtypes are unclustered and disordered (Hanley et al., 2016). These phenotypes resemble *Mnx1*-deficient mice, which form MNs in normal numbers yet show abnormal MN organization and axon elongation (Arber et al., 1999; Thaler et al., 1999). Due to the limitations of both transcriptomic and epigenomic techniques and the computational inference of DEGs, differential binding peaks and enhancer-gene pairs, we cannot rule out the possibility that additional direct targets of *Mnx1* have been excluded. However, our current results suggest that *Pou6f2* and particularly *Pbx3* are the two most likely direct targets of *Mnx1*. However, whether and how these two genes are regulated by *Mnx1*, and the rules of *Mnx1* in mediating the function of MNs *in vivo* remain to be explored.

Overall, this study represents a systematic investigation into the molecular function of *Mnx1* in MNs. Our results establish the binding preference, regulatory function, and putative gene targets of *Mnx1* in MNs, and identified *Pou6f2* and *Pbx3* as two putative direct targets of *Mnx1*. Furthermore, we observed unexpected properties of *Mnx1* as a transcription factor, which may be useful for the understanding the full regulatory potential of homeobox transcription factors.

## Methods

### Transgenic mouse ESC and CRISPR-Cas9 engineering

The DOX-inducible mouse ESC line was constructed by first generating a NIL transgene consisting of *Ngn2*, *Isl1*, and *Lhx3,* linked with 2A sequences, in the p2lox vector and performing the inducible cassette exchange reaction in A2lox ESCs as described (Iacovino et al., 2011). To generate the endogenous GFP tag at the *Mnx1* locus in A2lox NIL ES cells, we generated guide RNAs targeting a site close to the putative translation start site and cloned into the pX330 vector using BbsI (Addgene 42230), as summarized in **Table S3**. This CRISPR-Cas9 plasmid was co-electroporated into A2lox NIL ESCs using a Nucleofector (Lonza), with a repair plasmid containing the GFP coding sequence and flanking homology arms to generate an in-frame fusion of GFP with MNX1, along with an additional plasmid containing a hygromycin resistance gene. ESCs were treated with hygromycin for 48 hours before growing in regular ESC media at 37°C and 5% CO_2_ for one week. Single colonies were picked, expanded and screened by PCR.

To achieve exact deletion of desired *Mnx1* genomic fragments, two guide RNAs (gRNAs) and one single-stranded oligodeoxynucleotide (ssODN) were used per CRISPR experiment, with their sequences summarized in **Table S4**. gRNAs were cloned into the BbsI site of pX330. To construct the knockout cell lines, 10 μg of pX330 neomycin-resistant vector containing each gRNA and 10 mM ssODN were co-transfected into 2 × 10^6^ ESCs using a Nucleofector (Lonza, program A023). Cells were plated onto two 10 cm dishes containing hygromycin-resistant DR4 MEFs. Two dishes were selected with 150 μg/mL hygromycin for 48 hours and colonies picked one week later. Single clones were screened by PCR. The PCR primers used are listed in **Table S5**. Candidate knockout clones were further verified by sequencing of the purified PCR fragments.

### Mouse ESC culture and MN induction

Mouse ESCs were grown in ES medium consisting of DMEM (Gibco, 11965092) supplied with 15% FBS (HyClone, SH3007103), 1× HEPES (Gibco, 15630080), 1× NEAA (Gibco, 11140050), 1× GlutaMax-I (Gibco, 35050061), 1 ng/μL LIF (Gibco, A35933), 0.1% 2-Mercaptoethanol (Gibco, 21985023) and 1× Anti-Anti (Gibco, 15240062). Mouse ESCs were cultured on cell culture dishes a 37 °C with 5% CO_2_. The MN induction procedure takes 6 days. Day 0: mESCs were treated with trypsin to make a single-cells suspension, and ~5 × 10^6^ cells were seeded to 15-cm Petri dishes with 30 mL ADFNK medium consisting of 44% Advanced DMEM (Gibco, 12634010), 44% Neurobasal (Gibco 21103049), 10% Knockout serum replacement (Gibco, 10828028), 1× GlutaMax-I (Gibco, 35050061), 0.1% 2-Mercaptoethanol (Gibco, 21985023) and 1× Anti-Anti (Gibco, 15240062). Cultures were incubated on a rotating shaker at ~50 RPM at 37 °C with 5% CO_2_ for two days to generate embryoid bodies. Day 2: Embryoid bodies were collected by gravity, ADFNK medium was exchanged for ADFNK containing 1 μM RA (Sigma, R2625-100MG) and 1 μM SAG (EMD Millipore, 566660-5MG), and embryoid bodies were cultured with rotation for two days to generate neuronal progenitor cells. Day 4: Embryoid bodies were collected by gravity, medium was exchanged for ADFNK medium with 1 μM RA (R2625-100MG), 1 μM SAG (EMD Millipore, 566660-5MG), 5 μM DAPT (Sigma, D5942-5MG), and 1 μM DOX (Sigma, D9891-5G), and further culture for with rotation for two days to generate mature iMNs. Day 6: iMNs were collected, dissociated into single cells using the Papain Dissociation System (Worthington Biochemical, LK003150), and subjected to FACS using the BD FACSCalibur platform to evaluate the induction efficiency based on the GFP signal.

### MN axon elongation

MN axon elongation was performed as previously described (Wu et al., 2012) with modifications. In brief, iMNs were dissociated into single cells using the Papain Dissociation System (Worthington Biochemical, LK003150) after collection at Day 6 and resuspended to the concentration of 0.2 × 10^6^ cells/mL with MN medium consisting of Neurobasal medium (Gibco 21103049) with 1× B27 (Gibco, 17504044), 2% FBS (HyClone, SH3007103), 1× GlutaMax-I (Gibco, 35050061), 50 μM L-Glutamic acid (Sigma, G1251-100G), 10 pg/mL BDNF (R&D systems, 248-BD-005), 10 pg/mL GDNF (R&D Systems, 257-NT-010) and 1× Anti-Anti (Gibco, 15240062). The resuspended iMN cells were seeded onto coverslips within 24-well cell culture dishes, which were coated overnight with 10 μg/mL Poly-DL-Ornithine (Sigma, P0421) and 10 μg/mL Laminin (Millipore, CC095). Subsequently, the iMNs were cultured at 37 °C with 5% CO_2_ for two days (Day 8) and then used for further experiments.

### Western blotting

Cells were lysed at 4 °C for 2 hours in ice-cold lysis buffer (50 mM Tris-HCl (pH 8), 150 mM NaCl, 1% Triton X-100) supplemented with protease inhibitor cocktail (Roche, 11697498001). The supernatant was collected after centrifugation at 13000 RPM at 4°C for 20 min. Protein concentration was determined using the BCA protein assay (Thermo, 23225), and then 10 μg of protein per sample was mixed with SDS and reducing reagent and loaded onto NuPAGE Bis-Tris protein gels (Invitrogen, NP0322) after boiling at 70 °C for 10 min. Western blots were performed with the primary antibodies anti-GFP (Invitrogen, A11122), anti-MNX1 (gift from Samuel Pfaff)) and anti-GAPDH (Novus, NB300327). The secondary antibody was Goat anti-Rabbit IgG (Invitrogen, 656120), and signal detection was conducted using the Amersham ELC Select Western Blotting Detection Reagent (GE, RPN2235).

### Immunofluorescence

Immunofluorescence imaging was performed for iMNs either collected at Day 6 or after two days of axon elongation (Day 8). The samples were rinsed in PBS at RT for 5 min, fixed with 4% (v/v) paraformaldehyde in PBS at RT for 5 min, rinsed in PBS at RT for 5 min, washed in PBT (PBS with 0.2% v/v TritonX-100) at RT for 5 min three times, then blocked in 10% goat serum in PBST for 1 hour. Samples were incubated with primary antibodies at 4°C overnight and washed with PBST at RT for 5 min three times. Samples were then incubated with secondary antibodies in 1% serum in the dark at RT for 2 hours. The primary antibodies used included anti-GFP (Invitrogen, A11122, 1:2000), anti-Vsx2 (Chemicon, AB9016, 1:400) and anti-TUJ1 (Invitrogen, 480011, 1:250), and secondary antibodies included species-appropriate Alexa 488 (Invitrogen, A-21206, 1:500), Alexa 555 (Invitrogen, A21436, 1:500) and Alexa 647 (Invitrogen, A21235, 1:500). Samples were counterstained with Hoechst (Invitrogen, 33258, 1:10,000 in PBS) at RT for 5 min, rinsed with PBS at RT for 3 min twice, then mounted and dried in the dark. Finally, immunofluorescence imaging was performed using a Leica DM6000 B microscope.

### ChIP-Seq

ChIP-Seq was performed as previously described (Blecher-Gonen et al., 2013) with minor modifications. Chromatin fragmentation was performed using the Diagenode Bioruptor Plus sonicator. The antibodies used include: anti-GFP (Invitrogen, A11122), anti-H3K4me1 (Abcam, ab8895), anti-H3K4me3 (Millipore, #04-745), anti-H3K27ac (ActiveMotif, #39685), anti-H3K27me3 (Abcam, ab6002), and anti-H3K9me3 (Abcam, ab8898). The amount of chromatin used for each experiment was 30 μg for transcription factors and 20 μg for histone modifications. ChIP-Seq libraries were constructed using the Takara SMARTer ThruPLEX DNA-Seq Kit (R400674) and sequenced as 50 bp single-end reads with a HiSeq2500 instrument (Illumina).

Reads were trimmed using Trim Galore v0.6.1 and then aligned to the reference genome (mm10) using Bowtie v2.3.5 (Langmead and Salzberg, 2012) with default parameters. The PCR duplicates for each dataset were removed using the *rmdup* function of samtools v1.9 (Li et al., 2009). The data reproducibility between biological replicates was confirmed, then reads from replicates were pooled together for further analysis. Peak calling was performed with MACS v2.2.5 (Zhang et al., 2008) with parameters: −g mm −q 0.05. The called peaks were further cleaned by removing those that overlapped with ENCODE Blacklist V2 regions (Amemiya et al., 2019). Peaks from H3K27ac and GFP-MNX1 ChIP-Seq were used for further analysis.

### RNA-Seq

Total RNA was extracted using the RNeasy Micro kit (Qiagen) with on-column DNase digestion. RNA samples were submitted for library construction with the TruSeq stranded mRNA sample preparation kit (Illumina) and sequencing as 75 bp paired-end reads with a HiSeq2500 instrument (Illumina). Reads were trimmed using Trim Galore v0.6.1, and then aligned to the reference genome (mm10) using STAR v2.6.1 (Dobin et al., 2013). To identify differentially expressed genes, we first obtained the read count for each gene with the *featureCount* function of subread v1.6.3 (Liao et al., 2013), then identified differentially expressed genes with FDR<0.05 and fold change>2 using DESeq2 v1.22.2 (Love et al., 2014). The Transcripts Per Million (TPM) value for each gene was calculated using RSEM v1.3.2 (Li and Dewey, 2011). The tissue specificity index was calculated as previously described (Yanai et al., 2005).

### Motif enrichment analysis

Using the top 500 peaks in terms of peak score from each ChIP-Seq dataset, we performed motif discovery using MEME-ChIP (Machanick and Bailey, 2011) with parameters: −meme-minw 8 - meme-maxw 12. The flanking +/− 200 bp sequences from the summit were used for analysis.

### Genome annotation and gene ontology analysis

The reference genome (mm10) and gene annotations were obtained from the UCSC Genome Browser (Haeussler et al., 2019). We used the *annotatePeaks.pl* script from HOMER (Heinz et al., 2010) with default settings to annotate the genomic distribution of identified peaks. Gene ontology enrichment analyses for differential genes were performed using DAVID functional annotation tools (Huang da et al., 2009), and for genomic regions were performed with GREAT (McLean et al., 2010). PhastCons annotation was retrieved from the UCSC Genome Browser (Haeussler et al., 2019). Sequence conservation as measured by Jensen-Shannon Divergence (JSD) is calculated as previously described (Capra and Singh, 2007).

### Computational identification of *Mnx1* targets

We reasoned that *Mnx1*-regulated loci should be those with altered H3K27ac after disruption of *Mnx1*, and also that *Mnx1*-regulated genes should be those associated with *Mnx1*-regulated loci and show differential gene expression after disruption of *Mnx1*. Based on this rationale, we designed an unbiased procedure to computationally infer *Mnx1*-regulated cis-elements and genes by integrating MNX1 binding, histone modifications, and gene expression data (**Figure S5A**). First, differential H3K27ac loci between WT and *Mnx1*-mutant iMNs were identified using DiffBind (Ross-Innes et al., 2012), with summit set as 500. The identified H3K27ac loci are putative *Mnx1*-targeted cis-elements. Then, differential genes between WT and *Mnx1*-mutant iMNs were identified using DESeq2 (Love et al., 2014) as aforementioned. Finally, by comparing the list of differential genes against differential H3K27ac loci, the putative *Mnx1* target genes were identified as differential genes with at least one H3K27ac loci significantly changed at the same direction. Interestingly, only a few up-regulated genes with increased H3K27ac loci were identified, while no down-regulated genes with decreased H3K27ac loci could be identified.

### Statistical analysis and visualization

Statistical analyses were performed using the R language (Team, 2020). ChIP-Seq and RNA-Seq tracks were visualized using IGV (Thorvaldsdottir et al., 2013), and ChIP-Seq heatmaps were created using DeepTools (Ramirez et al., 2014). Heatmaps for gene expression were visualized with the R package pheatmap (https://cran.r-project.org/web/packages/pheatmap).

### Data deposition

All data reported in this study have been deposited in the Gene Expression Omnibus (GEO) database with accession GSE143161.

## Acknowledgement

This work was supported by the Intramural Research Program of the NICHD, NIH for TSM, and the Priority Academic Program Development of Jiangsu Higher Education Institutions (PAPD) for MAS. JD was supported by the NIH-Oxford-Cambridge Scholars Program. We thank Dr. Samuel Pfaff for kindly providing MNX1 antibody, the National Institute of Child Health and Human Development (NICHD) Molecular Genomics Core for high-throughput sequencing. This study utilized the computational resources of the NIH High-Performance Computing Biowulf cluster (https://hpc.nih.gov) and the Yangzhou University College of Veterinary Medicine High-Performance Computing cluster.

