## Supplemental Figures for "Homeobox transcription factor MNX1 is crucial for restraining the expression of pan-neuronal genes in motor neurons"

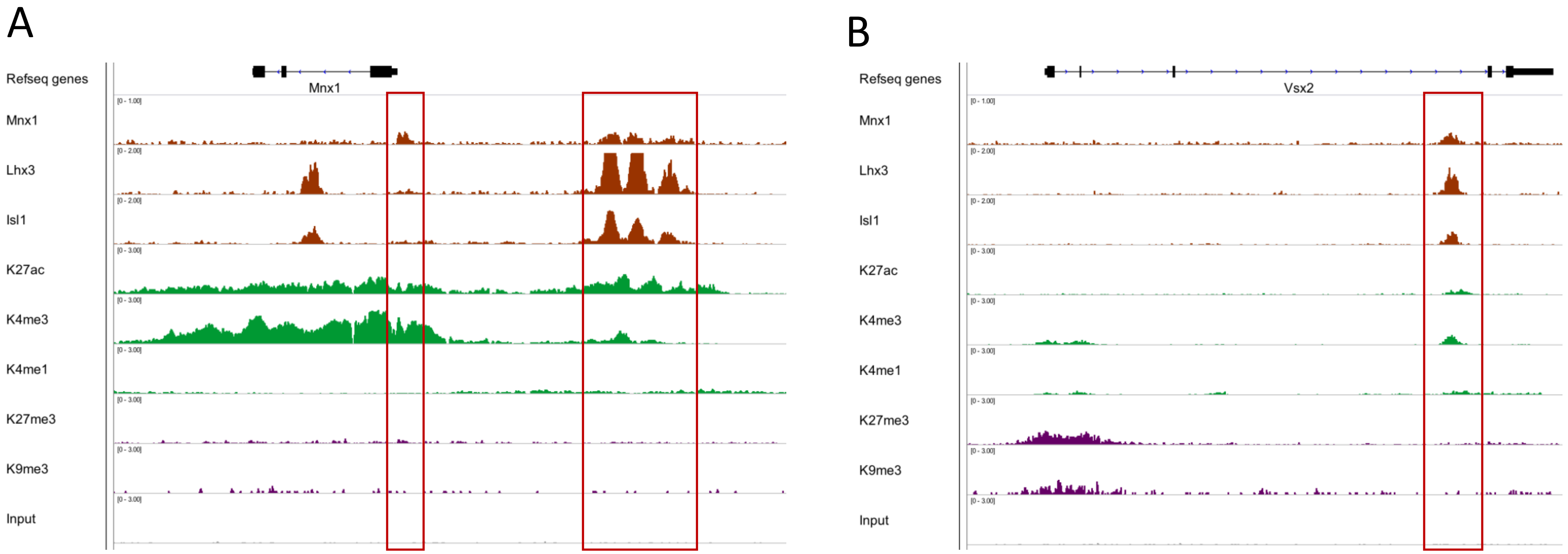


**Figure S1 MNX1 binding at *Mnx1* and *Vsx2* loci based on ChIP-Seq data**
The IGV tracks show MNX1 binding in *Mnx1* (A) and *Vsx2* (B) loci, which are known target genes for MNX1. Several histone marks are also shown. For *Mnx1*, MNX1 binding occurs in its promoter and up-stream regions. For *Vsx2*, MNX1 binding occurs in its 3$^{rd}$ intron. MNX1 bbound loci are highlighted with red rectangles.

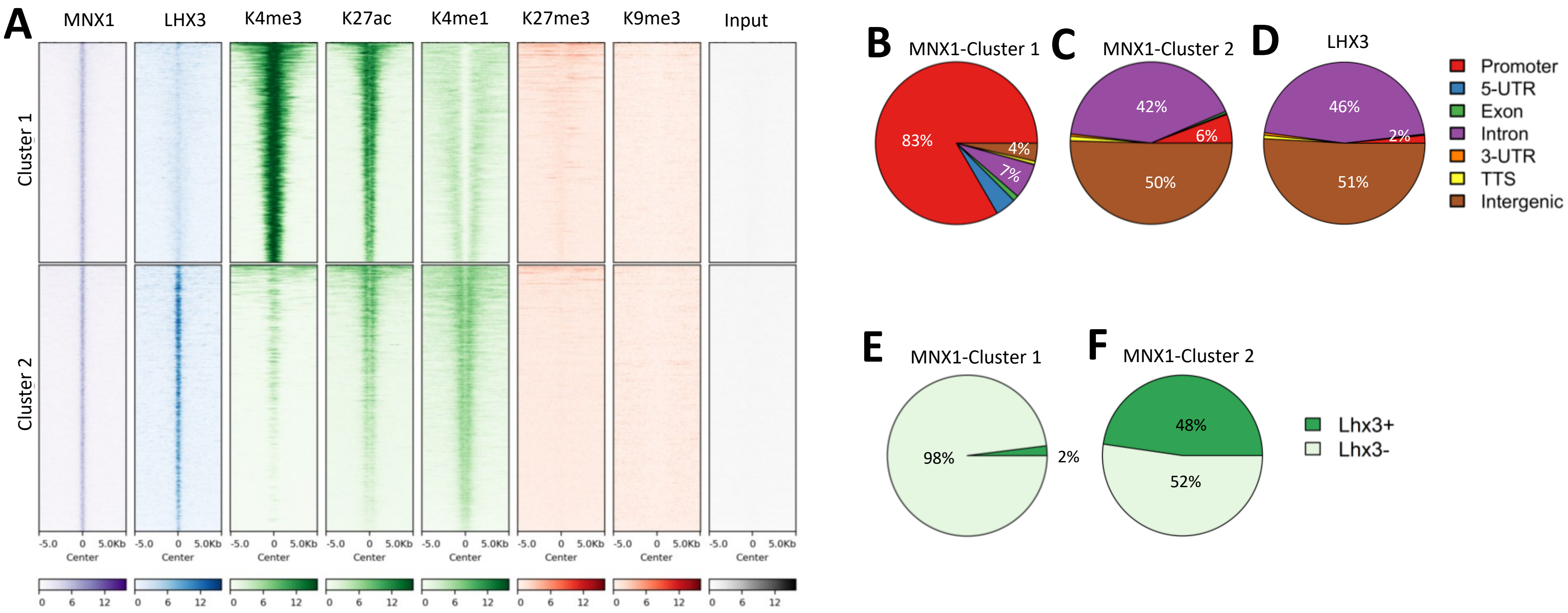


**Figure S2 MNX1 and LHX3 co-bind thousands of enhancers**

**A.** Heatmap showing the occupancy of MNX1, LHX3 and different histone marks for two groups of MNX1 binding loci. The two groups are generated using k-mean clustering (k=2) based on the occupancy of these factors. **B-D.** Pie plots show the genomic distribution of two groups of MNX1 bound loci and LHX3 peaks. **E-F**. Pie plots show the co-binding of LHX3 at two groups of MNX1 bound loci.

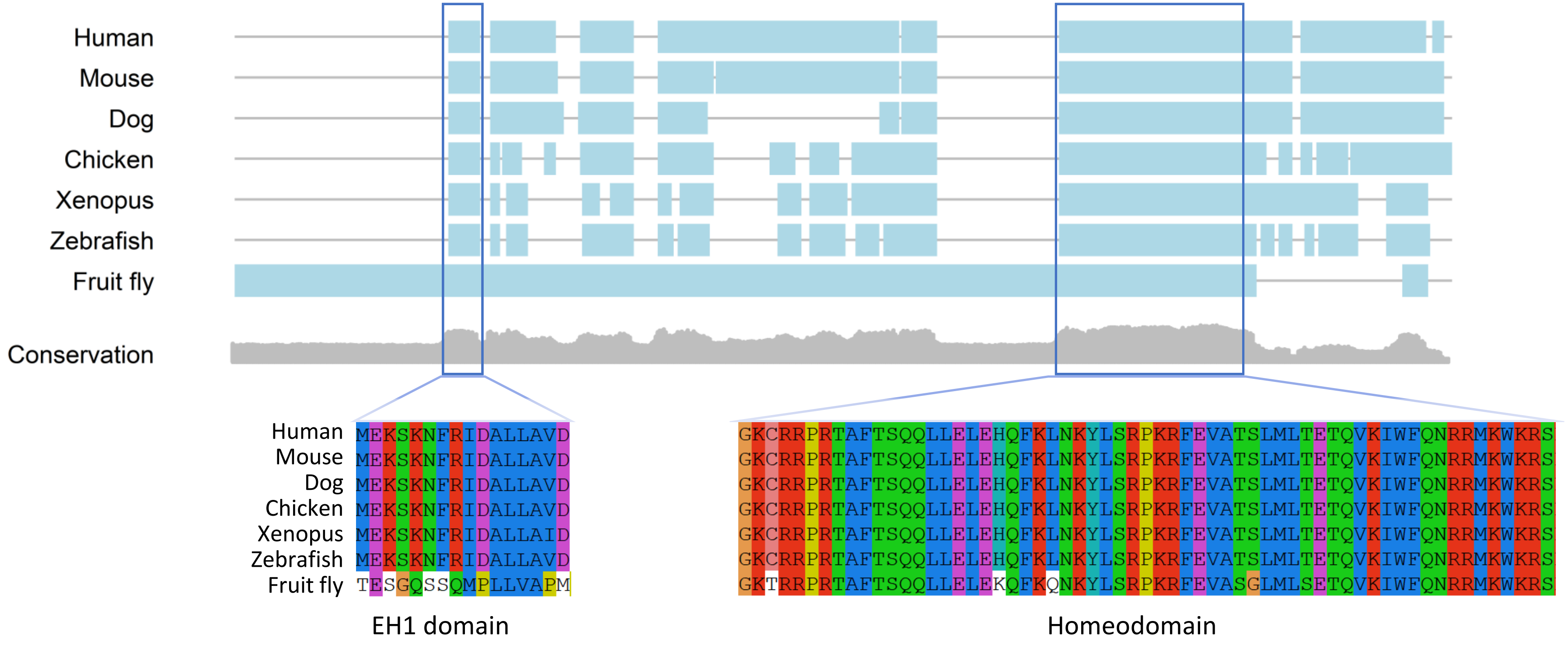


**Figure S3 *Mnx1* orthologous genes in different species share conserved EH1 motif and Homeodomain**

The top plot shows alignment of the protein sequences of *Mnx1* orthologues in 7 different species. Light blue rectangles indicate aligned sequences, while lines indicate gaps in the alignment. The middle plot shows sequence conservation along the alignment, as measured using Jensen-Shannon divergence (Capra et al., 2007). The bottom plots show the sequences of the aligned EH1 motif and Homeodomain in different species.

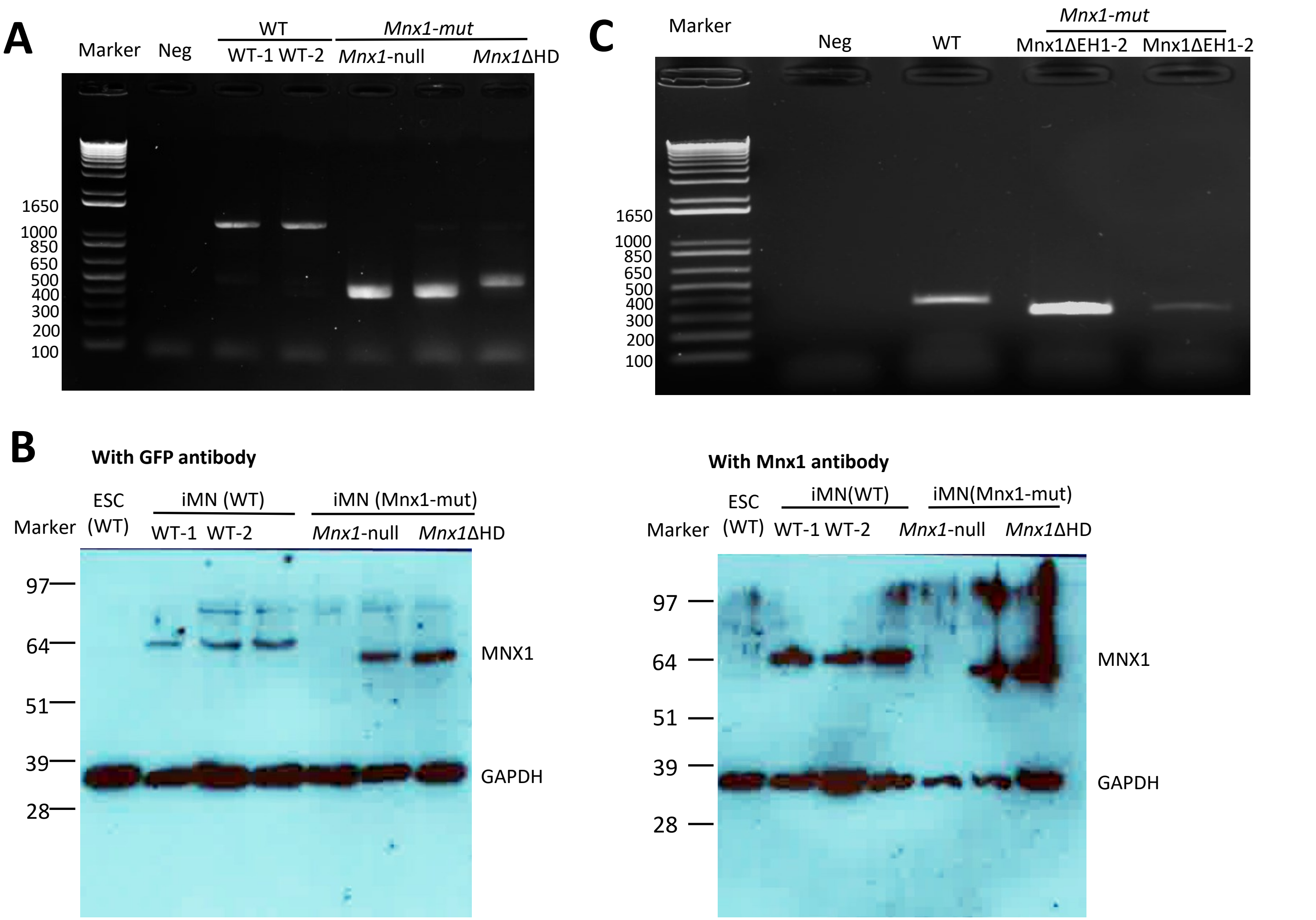


**Figure S4 Validation of different *Mnx1* mutants with deletion of EH1 motif and Homeodomain**

**A**. PCR results for the genotyping of WT, *Mnx*1ΔHD and *Mnx1*-null colonies. The marker (1 kb Plus DNA Ladder) and negative control are indicated. To be noted, the other replicate in the middle is not used in this study and is unlabeled here. **B**. Western blot showing the protein size and abundance for WT and ΔHD mutants. Protein sizes (kDa) based on ladder are indicated on the left. Protein from undifferentiated WT ESCs which do not express Mnx1 is used as negative control. Upper band is for MNX1, and bottom band is for GAPDH. Results using GFP (left) and MNX1 (right) antibodies are both shown. To be noted, while the lanes for three WT and three *Mnx1*ΔHD replicates are shown, only two replicates for each genotype are used in this study. The unused replicates are unlabeled here. **C**. PCR results for the genotyping of WT and *Mnx1*ΔEH colonies. The marker (1 kb Plus DNA Ladder) and negative control are indicated.

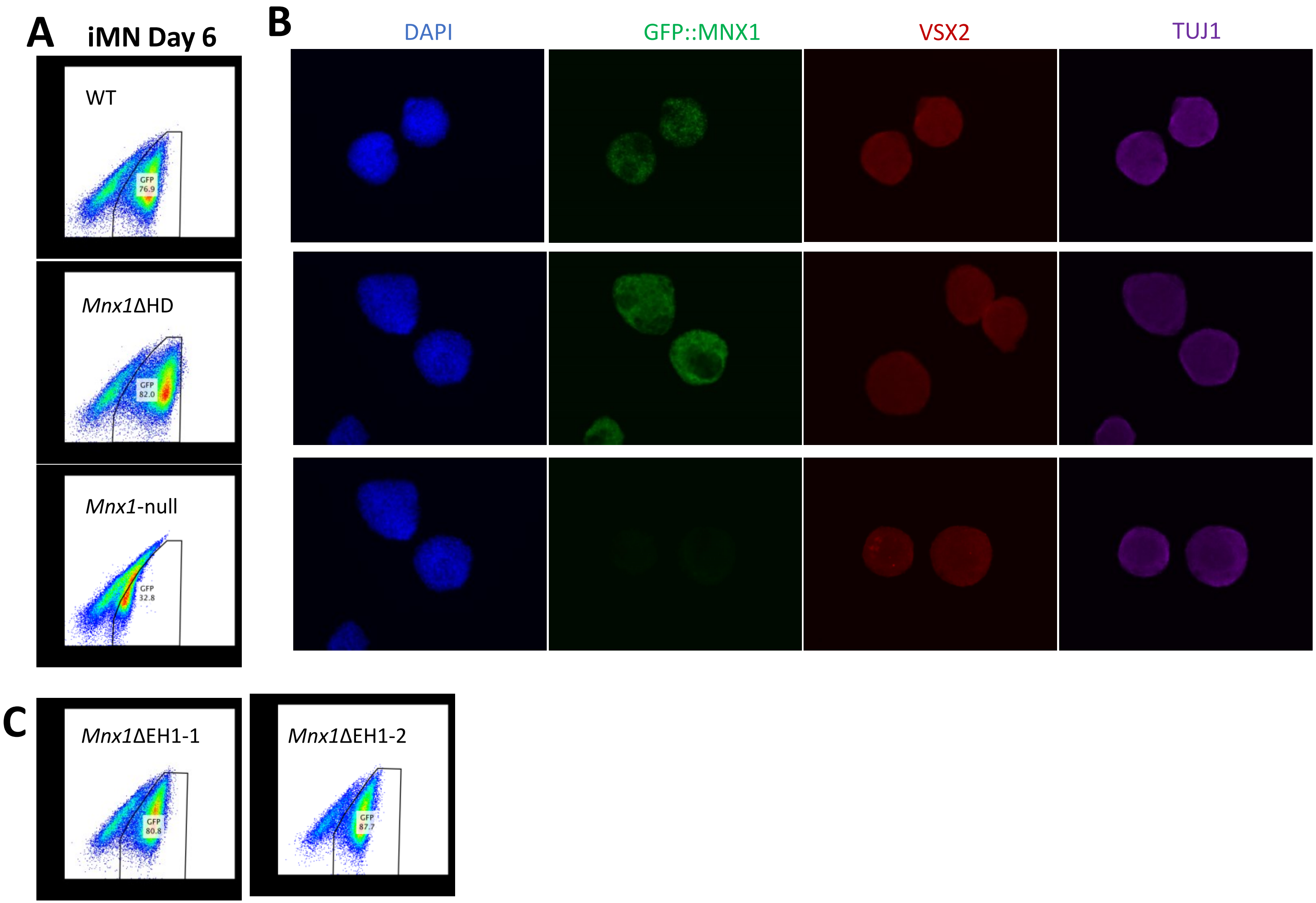


**Figure S5 Motor neurons can be induced at high efficiency for *Mnx1* mutants**

**A.** FACS sorting results for MNs collected at day 6 of directed induction. Results for WT and *Mnx1*ΔHD mutants are shown. To be noted, MNX1 protein in *Mnx1*-null iMNs is degraded, thus GFP::MNX1 signal is undetectable. **B.** Immunofluoresce imaging for embryoid bodies collected at day 6 of directed induction. Panels from left to right show the signals for DAPI, GFP::MNX1, VSX2 and TUJ1, respectively. Similar to A, *Mnx1*-null lacks GFP::MNX1 signal. **C.** FACS sorting results for iMNs collected at day 6 of directed induction. Results for wild-type and *Mnx1*ΔEH1 mutants are shown.

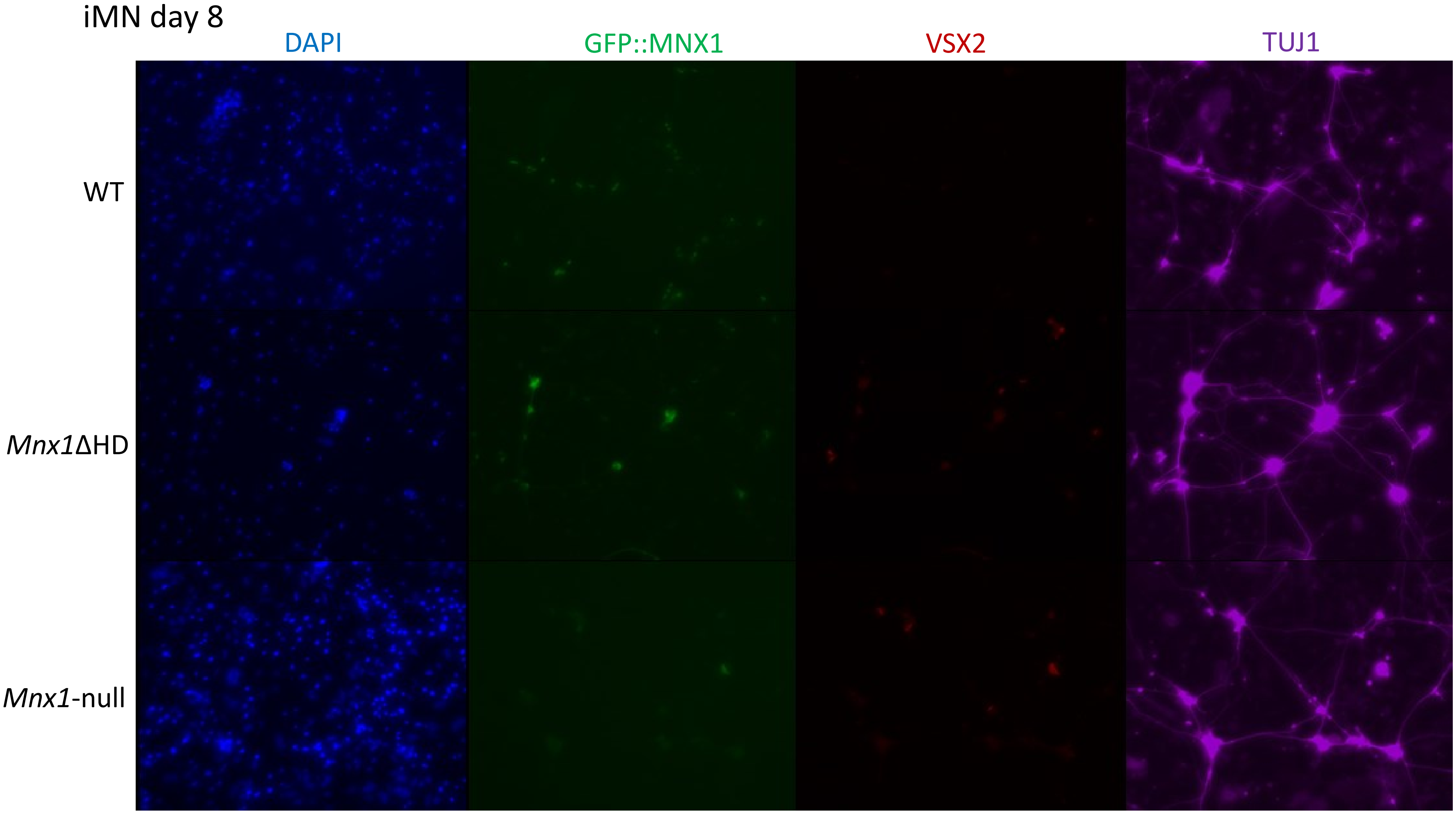


**Figure S6 *Mnx1* mutants have normal axon elongation under *in vitro* culturing**

This figure shows the immunofluorescent imaging for iMNs collected two days later (day 8) after 6-day induction. Panels from left to right show the signals for DAPI, GFP::MNX1, VSX2 and TUJ1, respectively. To be noted, *Mnx1*-null lacks GFP::MNX1 signal due to degradation of MNX1 protein.

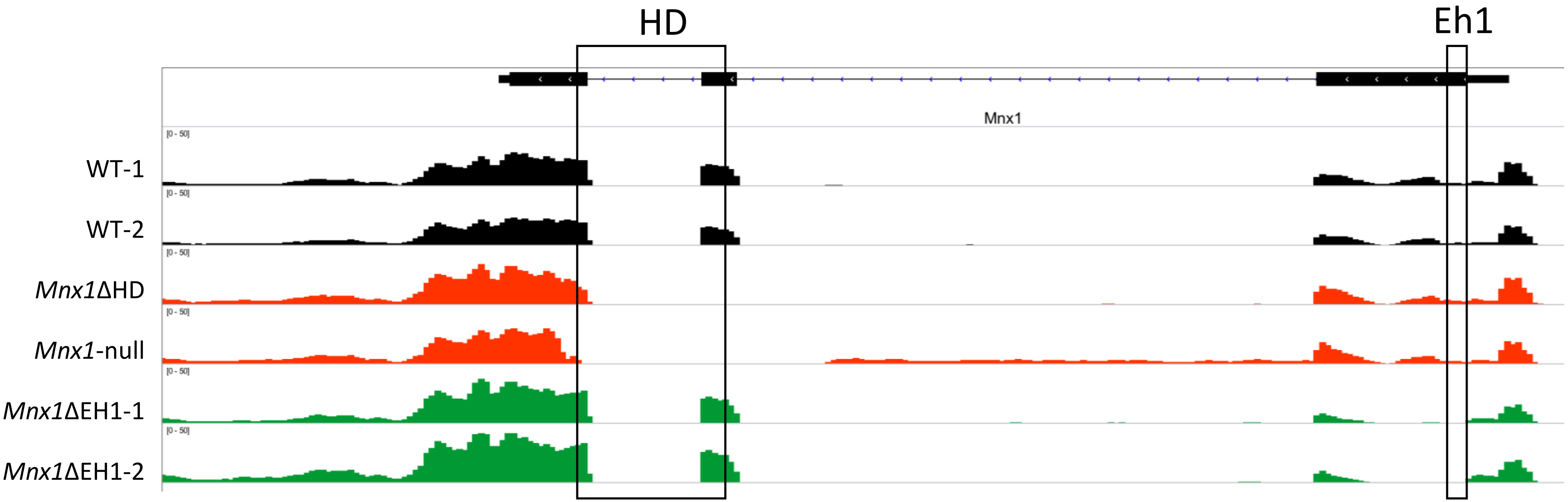


**Figure S7 Expression of *Mnx1* for different mutants and wild-type iMNs**
Normalized IGV tracks for the expression profile of Mnx1 among different *Mnx1* mutants and wild-type iMNs. The deletion of the homeodomain and Eh1 motif is evident for the corresponding samples. RNA abundance shows no remarkable difference between different mutants and wild-type iMNs. The positions for the homeobox domain and Eh1 motif are indicated by the rectangles.

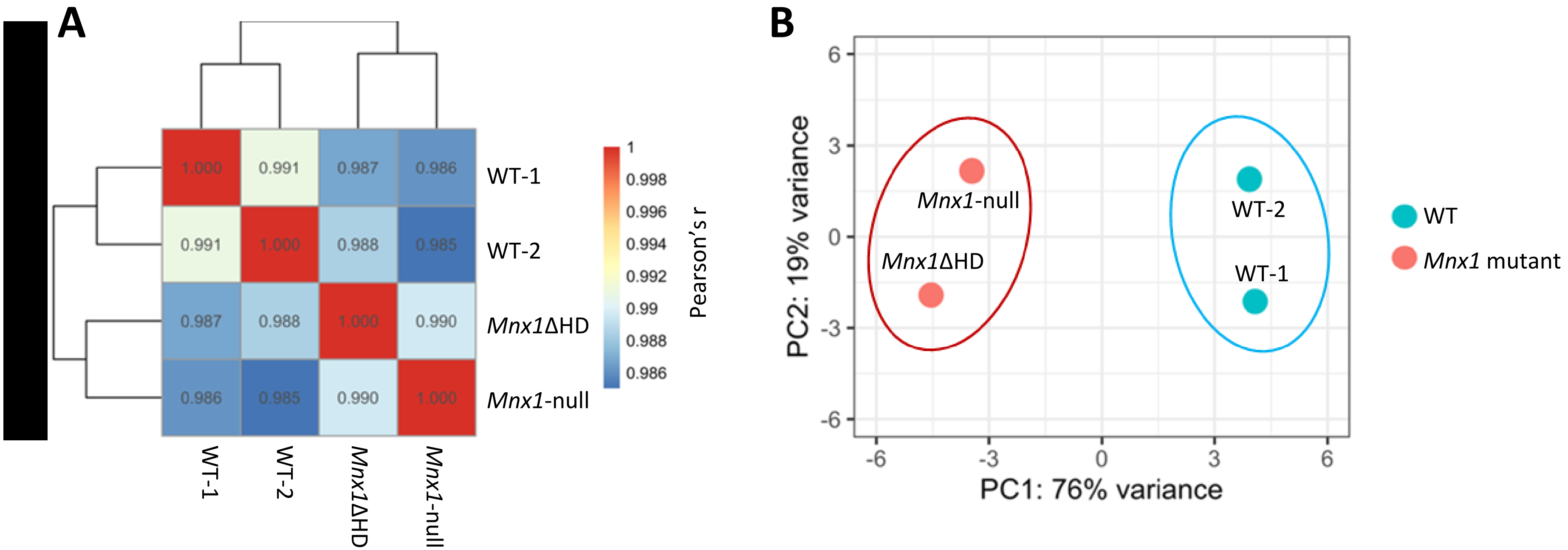


**Figure S8 Relationship among different WT and *Mnx1* mutants based on gene expression profile**
**A**. Heatmap showing the clustering of different wild-type and *Mnx1* mutant samples based on correlation among these samples. Color bar indicates Pearson's r, which is calculated based on log10(count) for all genes. Pairwise Pearson's r values are also marked in the heatmap. **B**. Relationship among wild-type and *Mnx1* mutant samples based on Principal Component Analysis. PCA is performed based on rlog-scaled counts calculated using DESeq2 package for the top 1000 genes with the highest variance among these samples.

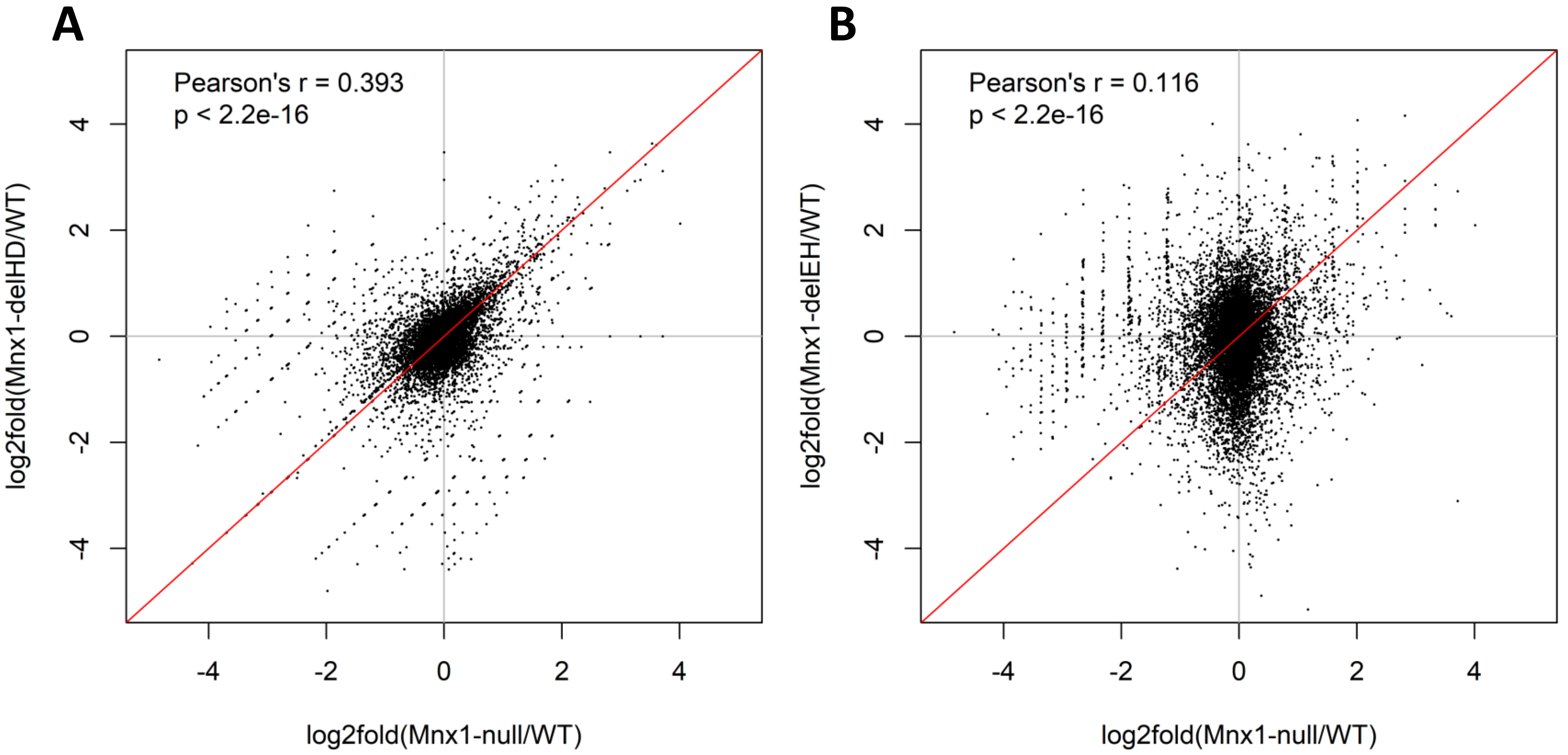


**Figure S9 Deletions of homeodomain and EH1 motif on gene expression relative to *Mnx1*-null mutant**
The scatter plots show the correlation of *Mnx1*ΔHD (A) and *Mnx1*ΔEH (B) on gene expression relative to *Mnx1*-null mutants. Pearson's correlation is calculated based on log2 fold changes of normalized read counts for all genes compared between *Mnx1* mutants and wild-type MNs.

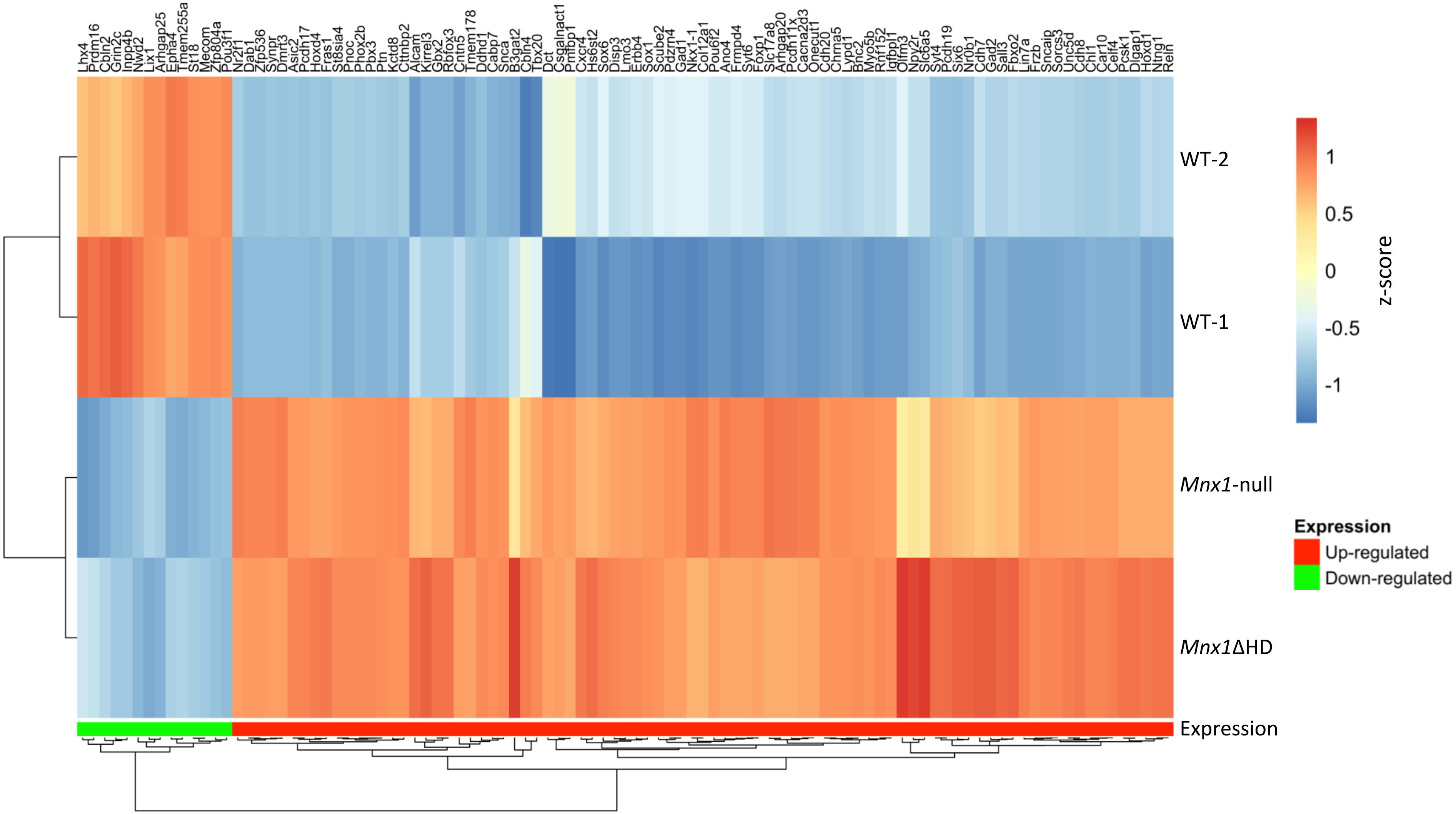


**Figure S10 Expression pattern of identified differential genes in wild-type and *Mnx1* mutant iMNs**

Heatmap of the expression pattern of the ninety-nine identified differential genes in wild-type and *Mnx1*ΔHD MNs. Gene names are labelled at the top. The groups of differential expression (up-regulated or down-regulated) are marked at the bottom. The color gradient indicates the *Z*-score.

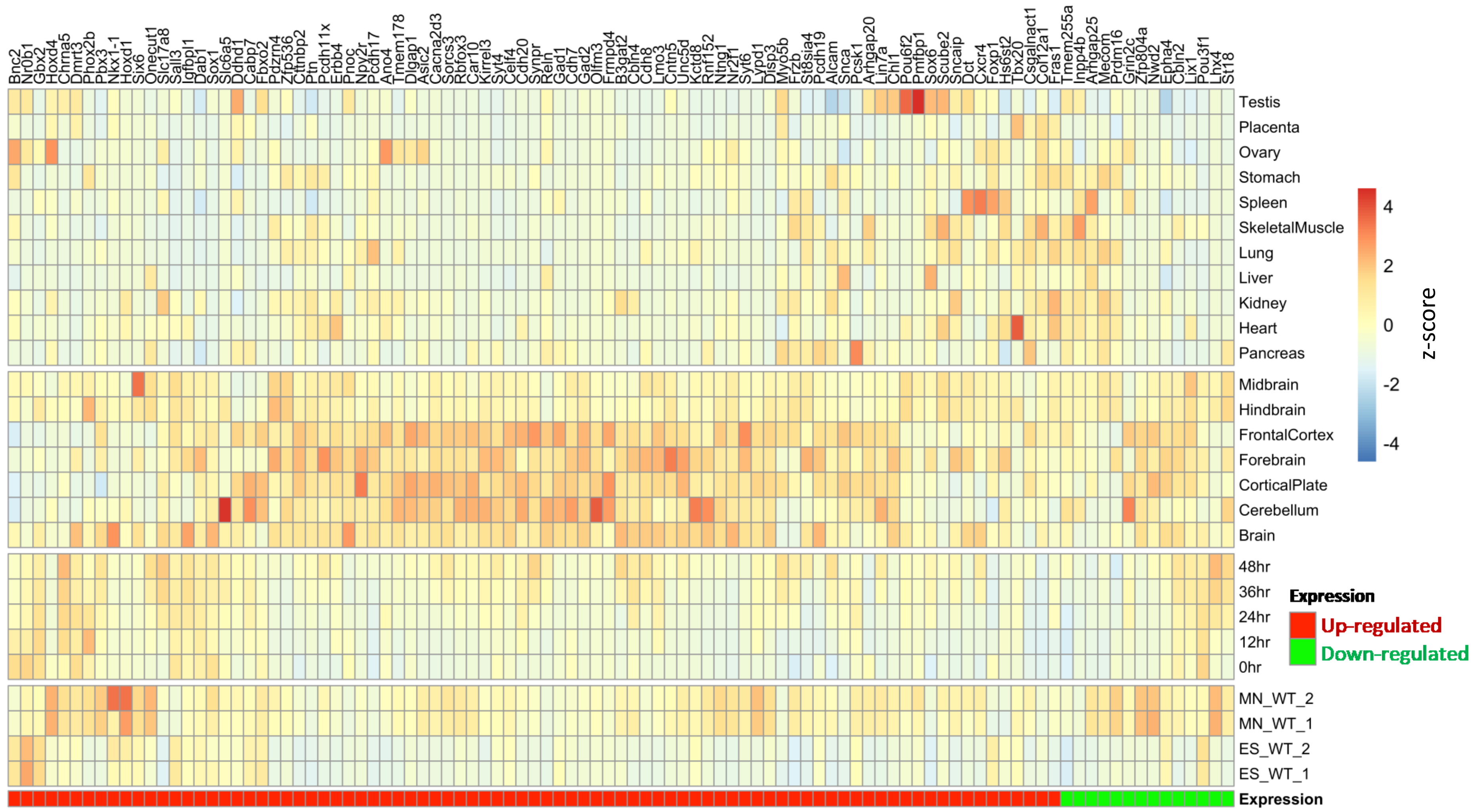


**Figure S11 Expression pattern of identified differential genes in different cell types and tissues.**
Heatmap of the expression pattern of the ninety-nine identified differential genes in ESCs, iMNs and different tissues. The groups of DEGs (up- or down-regulated) are marked at the bottom. The color gradient indicates the Z-score.

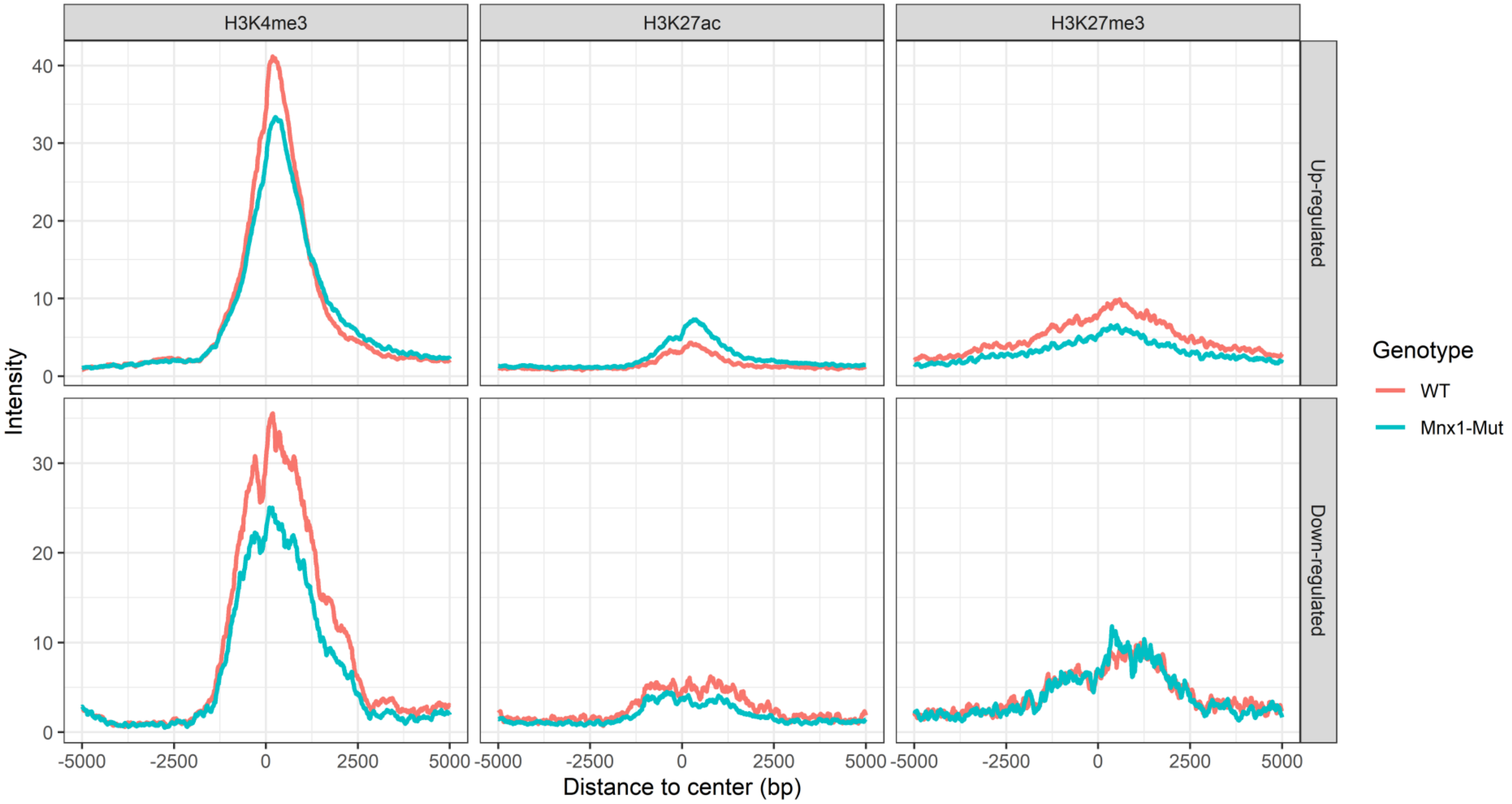


**Figure S12 The alteration of the occupancy of histone marks for the differential genes after *Mnx1* knockout.**
The averaged curves show the occupancy of different histone marks flanking the promoters of differentially expressed genes between WT and *Mnx1* mutant iMNs.

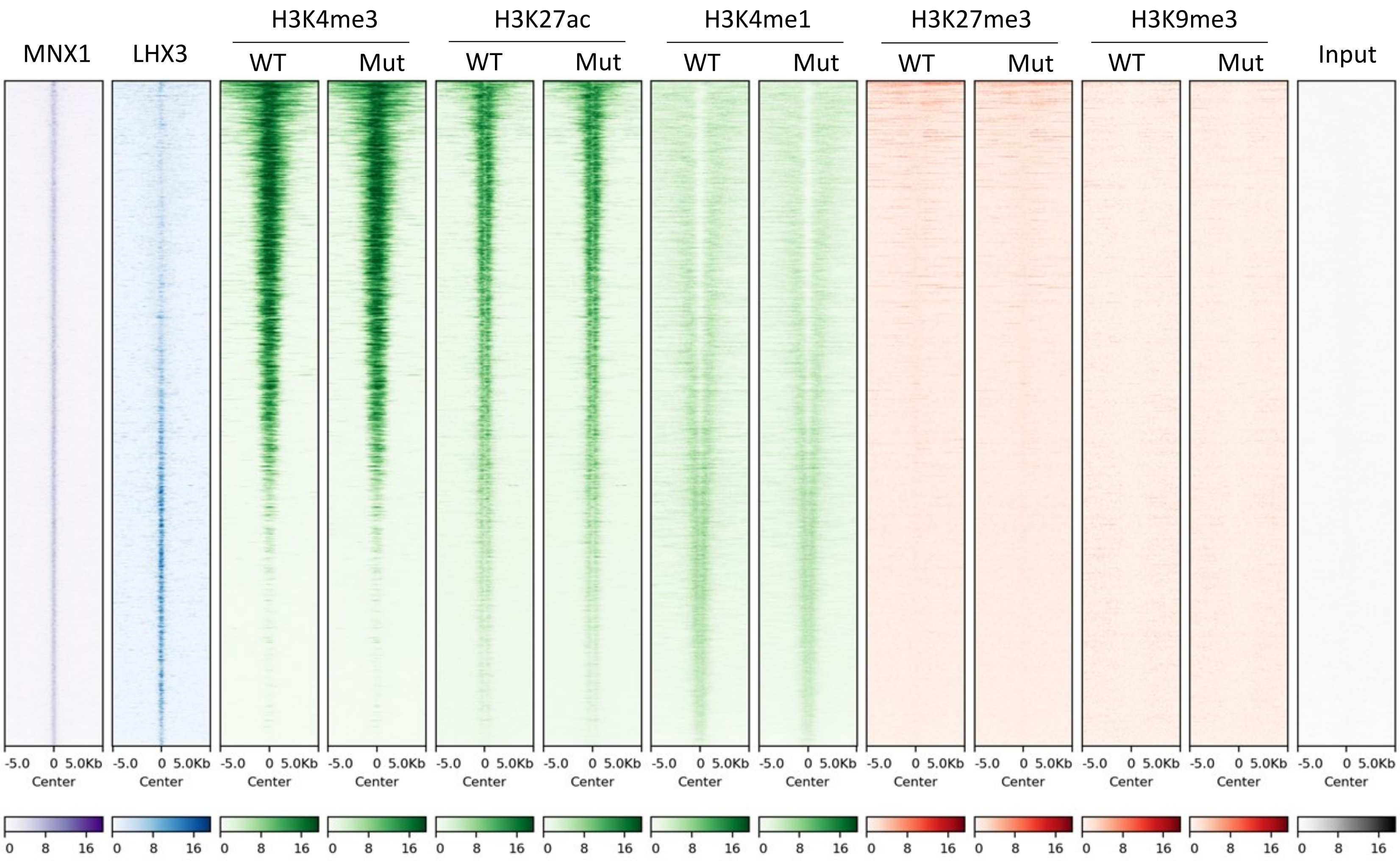


**Figure S13 Disruption of *Mnx1* has no global effects for the histone modification patterns on MNX1 bound loci**
The heatmaps show the occupancy of MNX1, LHX3 and different histone marks (H3K4me3, H3K27ac, H3K4me1, H3K27me3 and H3K9me3) flanking MNX1 bound loci in WT and *Mnx1*-mutant (ie. *Mnx1*ΔHD and *Mnx1*-null) iMNs. The color indicates the binding intensity calculated from ChIP-Seq data.

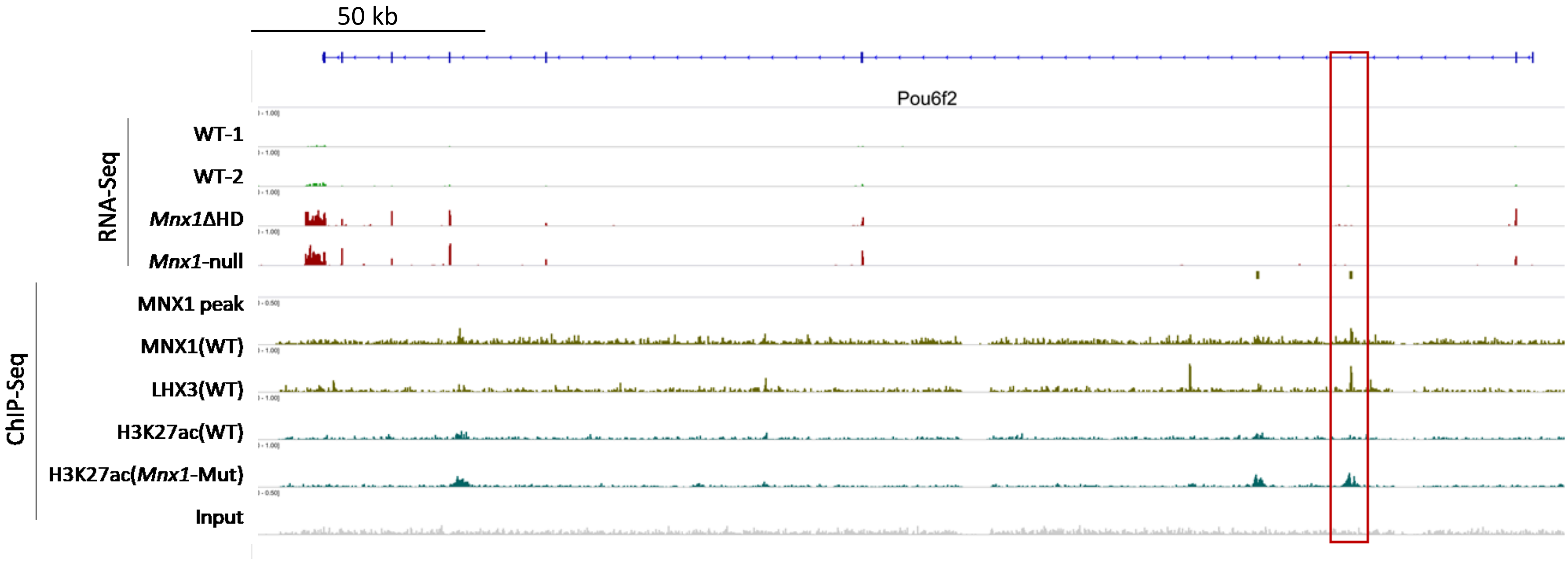


**Figure S14 Differential expression and occupancy of different factors on Pou6f2 locus.**
The IGV tracks show the differential expression and occupancy of MNX1, LHX3 and H3K27ac on Pou6f2 locus for WT and *Mnx1*ΔHD iMNs. The putative Mnx1-regulated loci is highlighted by red rectangle.

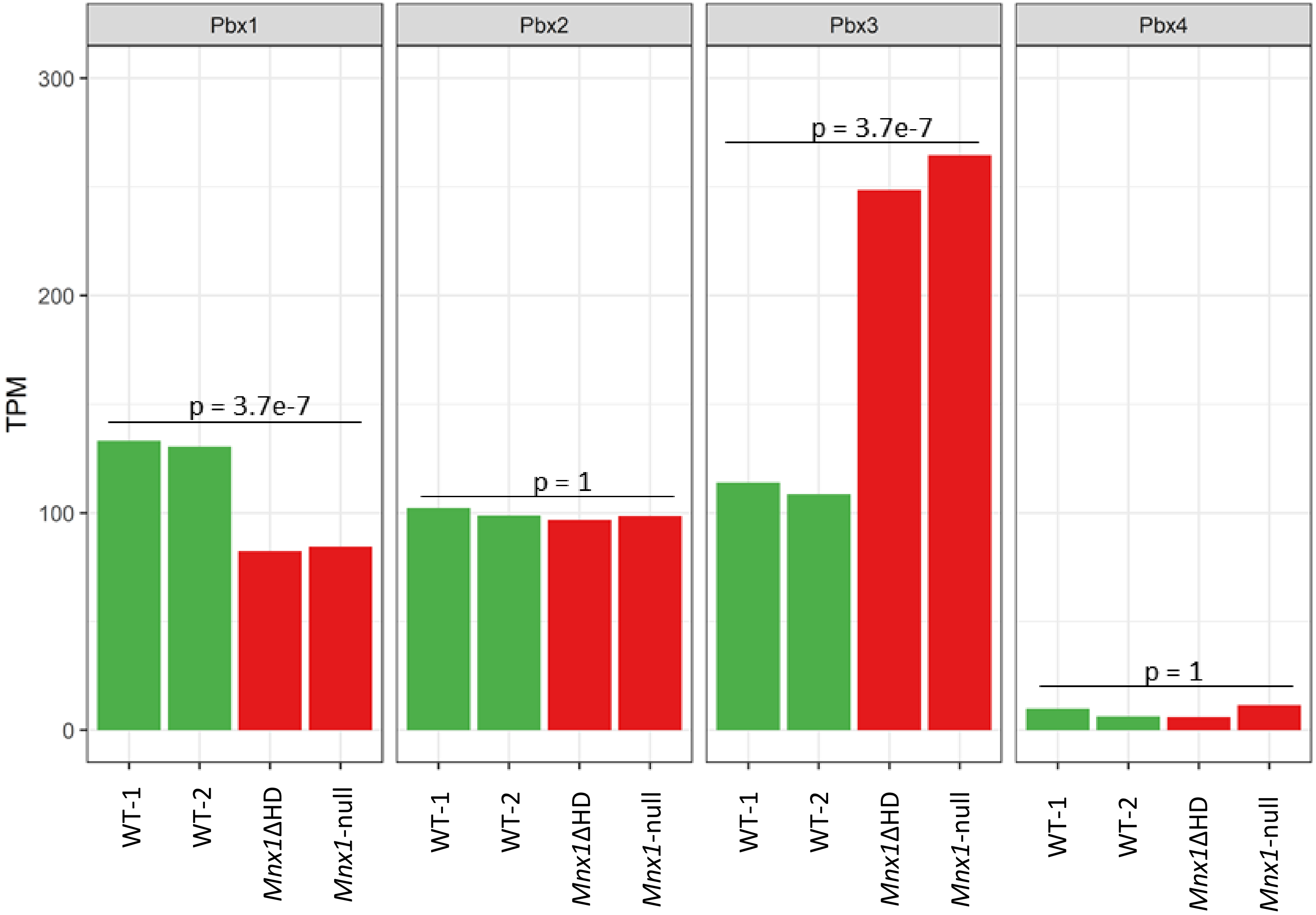


**Figure S15 Differential expression of *Pbx* genes in iMNs after disruption of *Mnx1***
This figure shows the expression values of the four *Pbx* genes (*Pbx*1-4) in WT and *Mnx1* mutant (ie. *Mnx1*ΔHD and *Mnx1*-null) iMNs. The y-axis indicates TPM values. P-values from DESeq2 results are indicated for each gene.
