## Supplementary material for "Homeobox transcription factor MNX1 is crucial for restraining the expression of pan-neuronal genes in motor neurons": Table S3

**Table S3 The pX330 plasmid information and sequence**

| Feature Name | Start | End |
|---|---|---|
| pBR322_origin | 168 | 787 |
| Amp | 942 | 1802 |
| Infusion HD extra 4bp | 2420 | 2423 |
| gRNA homology mutation 1 | 6207 | 6207 |
| gRNA homology mutation 2 | 6209 | 6209 |
| gRNA homology mutation 3 | 6211 | 6211 |
| 5'Arm homology | 2424 | 6214 |
| EGFP | 6215 | 6931 |
| 3'Arm homology | 6932 | 10388 |

**pX330 Sequence:**

tcttccgcttcctcgctcactgactcgctgcgctcggtcgttcggctgcggcgagcggtatcagctcactcaaaggcggtaatacggttatccacagaatcaggggataac

gcaggaaagaacatgtgagcaaaaggccagcaaaaggccaggaaccgtaaaaaggccgcgttgctggcgtttttccataggctccgcccccctgacgagcatcacaa

aaatcgacgctcaagtcagaggtggcgaaacccgacaggactataaagataccaggcgtttccccctggaagctccctcgtgcgctctcctgttccgaccctgccgcttac

cggatacctgtccgcctttctcccttcgggaagcgtggcgctttctcatagctcacgctgtaggtatctcagttcggtgtaggtcgttcgctccaagctgggctgtgtgcacga

accccccgttcagcccgaccgctgcgccttatccggtaactatcgtcttgagtccaacccggtaagacacgacttatcgccactggcagcagccactggtaacaggattag

cagagcgaggtatgtaggcggtgctacagagttcttgaagtggtggcctaactacggctacactagaagaacagtatttggtatctgcgctctgctgaagccagttacctt

cggaaaaagagttggtagctcttgatccggcaaacaaaccaccgctggtagcggtggtttttttgtttgcaagcagcagattacgcgcagaaaaaaaggatctcaagaag

atcctttgatcttttctacggggtctgacgctcagtggaacgaaaactcacgttaagggattttggtcatgagattatcaaaaaggatcttcacctagatccttttaaattaaaa

atgaagttttaaatcaatctaaagtatatatgagtaaacttggtctgacagttaccaatgcttaatcagtgaggcacctatctcagcgatctgtctatttcgttcatccatagttgc

ctgactccccgtcgtgtagataactacgatacgggagggcttaccatctggccccagtgctgcaatgataccgcgagacccacgctcaccggctccagatttatcagcaat

aaaccagccagccggaagggccgagcgcagaagtggtcctgcaactttatccgcctccatccagtctattaattgttgccgggaagctagagtaagtagttcgccagtta

atagtttgcgcaacgttgttgccattgctacaggcatcgtggtgtcacgctcgtcgtttggtatggcttcattcagctccggttcccaacgatcaaggcgagttacatgatccc

ccatgttgtgcaaaaaagcggttagctccttcggtcctccgatcgttgtcagaagtaagttggccgcagtgttatcactcatggttatggcagcactgcataattctcttactgt

catgccatccgtaagatgcttttctgtgactggtgagtactcaaccaagtcattctgagaatagtgtatgcggcgaccgagttgctcttgcccggcgtcaatacgggataata

ccgcgccacatagcagaactttaaaagtgctcatcattggaaaacgttcttcggggcgaaaactctcaaggatcttaccgctgttgagatccagttcgatgtaacccactcgt

gcacccaactgatcttcagcatcttttactttcaccagcgtttctgggtgagcaaaaacaggaaggcaaaatgccgcaaaaaagggaataagggcgacacggaaatgttg

aatactcatactcttcctttttcaatattattgaagcatttatcagggttattgtctcatgagcggatacatatttgaatgtatttagaaaaataaacaaataggggttccgcgcac

atttccccgaaaagtgccacctgacgtctaagaaaccattattatcatgacattaacctataaaaataggcgtatcacgaggccctttcgtctcgcgcgtttcggtgatgacgg

tgaaaacctctgacacatgcagctcccggagacggtcacagcttgtctgtaagcggatgccgggagcagacaagcccgtcagggcgcgtcagcgggtgttggcgggt

gtcggggctggcttaactatgcggcatcagagcagattgtactgagagtgcaccatatgcggtgtgaaataccgcacagatgcgtaaggagaaaataccgcatcaggc

gccattcgccattcaggctgcgcaactgttgggaagggcgatcggtgcgggcctcttcgctattacgccagctggcgaaagggggatgtgctgcaaggcgattaagttg

ggtaacgccagggttttcccagtcacgacgttgtaaaacgacggccagtgaattcgagctcggtacccggggatcGCCATAAGAGGGAATCCGATTTCCTG

TAGCTGGAGTTGGAGGTGGTTGTGAGTGTGAGTGGTGGGAACCCAACTTGGGTCCTCCTCAAGAGCTGCACTCACTCTTAAGCA

CTGAACTGTGGAACCACTATACTCCCCACCCCAACATCTTATTTCCTTGTGGTTATTTGTATAAATCAACACACCTTTCTCAGTAAT

GAGAGGAGGCTCAATCACAAGTTATATTCAGTAACTTAATGCAAGACCCAAAAGAGGGATCCTGGCTAGACGCAGAGAAAAAA

ATGACTGGGAGCCAGATGATTAAATGTACACCGGGCCTTAAATTCCAGCCTCTTTTGGGTAGGAGAGGTCTTCCTCTCTGCCCTC

CAAGAGCCTCATTTGGCCCCTGGGGCTTCTGGAGAAGGGAGCTACCTTGAGGCACAACAATGGGAATGCTGTGCTCACTTCGG

CAGCACCTATACTAAAACTGAACCGATACAGAGAAGATTAGCATGGCCCCTGCGCAAGGATGACATGCAAATTTGTGAAGCATT

CCTTATTTTTTTTTGCCAGGACCAGGAATGGGAACAGGTGGGTTGGGGAGCAGAGGGAGGGGAGAGGGGATAGGGGATTTGG

GGAGGGGAAACTAGGAAAGGGGATAACATTTGAAATGTAAATAAAGAAAATATCTAATAAAAAAAAAAAAAAGAATAGGCATG

CTTCTGGGACCAGACCTTTGATCTCTCAAGGGTACCGTAGTAAACCCCCAGCTCAGTCCCAGGCTGCCCCAGAGCCAGCTGGG

CTGGGTCAATAGACGCTGGGAACAGGAAGAGACGCAGCATGCTCACTCCACTCGTAAAACACACAAAGATCCCAGTTCCAGAC
GACTCCCAGTCTCTCTCCCAACTCCCACCTCTCAGGATACAAGTTCCCGCTCAAACCGAAGCAGAATATATGATCATATTCAAGC
CCTTTGACTTCAGAGTCTTTTTGGCCCAACCCCCACTTTGTCAAACAAAGAAATATATCCATCAAGTCTGATATGCAAATTTCCCC
TTCGCTGAAGGGGCTCTTGATACAACCTCACTCAGCCCACAGGTGGTGGGGACTGGTGTGGGTTAGCCTGGTGGATGGTCAAA
CACCAGACGAAGTGGGAAGGGTTTGAAAGGGATTTAGAAGCTTTTGGATCACATCCCCTACACAACAAACAGGAAATGAATTC
CTCTTCTCGGATGACTTTACTAAGGAAACCTGTGCCTAAACTCAGTAGACCTAATACTGTATCATCACCAAAAACAACCCTGGTT
GTGTAGTTTAGAGATGATTCGGGTCCATGAGGTCCACCCCAGGAAAATCTCATCTGCAGCCACCCTTTTTTTCCTCCTTCTTCTCC
TTTGTTTGTTTTGTTGTTGTTTTTTTCAAGACAGGGTATCTCTGTGTAGCATTTTTGGTTGCTGTCCTGGAACTGGCTCTGTAGACTA
GGCTGGCCTAGAACTCACAGACCCCACCTGCCTCTGCCTCCTGAGTGCTGGGATTAGAGACTTGCCTCACCATGGTTAGGTTCT
CTTTCTTCTTTGGGACTCTCATCTGGAAAAGGCCCATGGGCCTTAAACACAAGTTTCTGTAAATGCAATGGGGTCAAGACTGAAT
CCTGGGGATGGGGGTAGGGCATCCCAGCATCCTTACTTGCCTTAATTGGGCTTCCAAGATAGGAGAACTGGCAGGGAACCACT
CATGGAACACAAGTGTTCCTTTAACCCTCTAAAAGGCAGAGCTGCATTTTCATCACCCCAAAGAGACAGCACCTAAAGACAAGA
GCTATGTGCTTGTACTATTTTCAGACTAGCTGGAGGCCTTTGCAAAGCTGTGCACAACTTACTCCAAGATGTTCCCCGCTGGGTT
CCAGATTTTCCAGCCAGGACCACCCTGTCATAATAAAAACTCCACTCAGCCGAGCATGGTGGCGCACACCTTTAATCCCAGCAC
TTGGGAGGCAGAGGCAGGAGGATTTCTGAGTTCGAGGCCAGCCTGGTCTACAGAGTGAGTTCCAAGACAGCCAGGGCTAGAC
AGAGAAACCCTGTCTCCAAAATCAAAAACAAAAACAAAACAAAAACAACAAAAACAAAAAACCTACTCACTCATCCCAACAGA
ATCACTGCGTTAAATCTCTCCTAAACAAAACTTGTTGACAGGGAAAATATAGTCCTTACTTATATTTATCGTTACCCCCCACCCCC
AATCCCTCTTGAAACAGCTTTGCCTGCTACTTTTAAATTCACCCAACAAGGATTGTCCTAGCCTCCTGGCCTTCCCTGGGCCATTT
CCGATCTTTTATTAACCTCTCACCTGGTTGGGAAGCCGCGATCCAGAGACCTGGAACTCCCTCCCACTTTCCCCTCGTGTTTGGG
GAAGATACCCGAGAATGGAGCTAGCGAGAAAGCTTGATTTCTCAAACTACTTATTGATTCGTTACATTAAAAAAAAAAAGCCCC
CGGAAGCGTGTTTACAGTAAATATTTGATATGTGCGCAGGCTGAGGTCTGGATTCACACTTTAATACGGTCTCTGGAAGCCCTTT
CTTGATTTTCCTGAACTCGGCCGGCAGGAGGAGGCGAGAGCCTGATAGTGGCTCCCTTAGCCTGCGAACTGGCTCAGTCCGAG
GGAGCAGTCCTCGGCCAATTCCCGCAAAGCCTTTTTCCTACACCCCCACAAGGCAAGGCTCTAATAGAACTTGGTTCTCACCTA
GGTCCGGCTCAGTGGCACCTGTGCAAACGACCTTCAAGTCAATGTGCATCTCTGAGTTGGAAAACTGCTCTCTCTCCACTCTGAC
TGAAGATTTGTAGGCTTCAACCTCTAATACCGTATTTATACACACCGAAACCCTACAAACAAACATTTGCGGCGCATGATACTTTT
GGGGGAGGGGCAGATAAGAAGGTTCGACCTGCTGAGGGTTAATGACAGGGACGCAAACCATCTTGAAGCAATCAACACTCAG
GAACGCGAGAGATTTTGGGAAACCGAACAGTCTTCAGTGTACTAAGCTCAGGGTCCCGTTTTGGGTACCTAGTGGACTTCAGGG
GTGAGGACCCACCTAGTTCCTCAGTGCAGAACATTCCACAACTCCCTCAAGGGAGCGCGAGTAGCTGGAGGAAGATAAGGCGT
CCGCCTCTGGAGGAACAAGGTGTCAAAGGGACTCGGGGGGCACTCACTGGGACCCAGGGTTACTGGCGCTCACCACTCTCCCA
GCAGCAAGGAAGCAGTAAGCTTGTGTTCAGCCTCCAAACTCTCCAACCTGCCTCCTCCGGCCACCTACGCTGTTCACCACCGGC
TGCTTAGGACCTGGGGGCGCAGGGGCGGGGCGTCAGGCTAATTGGACCGTGACAGGTGAGCCCCTCGGCCAATCGCCGGGCA
GACAGCCCTCCTCGGGCCCCCGCTGAGCGCGCCAATCCCAGACCGCCCTGAGCTTCAAGTGTGCGGAGGGGCGCGGGTCCCC
ACCAAGGCACCAGTGGCCTCGATGCGGTCGCTTGAGCTCTTGAGAGCAGAGAGCCGGTCTCCCCAACACAAACTCGCAGGAG
CGCTCCGGTGGAAGTTCATACCAGTGAGTGGAGCGCGGTCCCCACGAGGACCGCAAAGAGGAGGCGCCTGGACGATGGTCGC
TCTCAGCTGAGTTTCCGGCTGCGACTTTATTGGCAAAAAATCGCAGTAACAATACCGGCCCCAAGGCTGGCCACCGCCACGCTC
AGTCTCCGTAAGCCCGCACACGACTGCATCGCCCACCCTTGGACCCTGATTGTCAGCTCCAAcCtAtCCGATGGTGAGCAAGGG
CGAGGAGCTGTTCACCGGGGTGGTGCCCATCCTGGTCGAGCTGGACGGCGACGTAAACGGCCACAAGTTCAGCGTGTCCGGC
GAGGGCGAGGGCGATGCCACCTACGGCAAGCTGACCCTGAAGTTCATCTGCACCACCGGCAAGCTGCCCGTGCCCTGGCCCA
CCCTCGTGACCACCCTGACCTACGGCGTGCAGTGCTTCAGCCGCTACCCCGACCACATGAAGCAGCACGACTTCTTCAAGTCCG
CCATGCCCGAAGGCTACGTCCAGGAGCGCACCATCTTCTTCAAGGACGACGGCAACTACAAGACCCGCGCCGAGGTGAAGTTC
GAGGGCGACACCCTGGTGAACCGCATCGAGCTGAAGGGCATCGACTTCAAGGAGGACGGCAACATCCTGGGGCACAAGCTGG
AGTACAACTACAACAGCCACAACGTCTATATCATGGCCGACAAGCAGAAGAACGGCATCAAGGTGAACTTCAAGATCCGCCAC
AACATCGAGGACGGCAGCGTGCAGCTCGCCGACCACTACCAGCAGAACACCCCCATCGGCGACGGCCCCGTGCTGCTGCCCG
ACAACCACTACCTGAgCACCCAGTCCGCCCTGAgCAAAGACCCCAACGAGAAGCGCGATCACATGGTCCTGCTGGAGTTCGTG

ACCGCCGCCGGGATCACTCTCGGCATGGACgAgCTGTACAAGGAAAAATCCAAAAATTTCCGCATCGACGCCCTGCTGGCCGT
GGATCCCCCGCGAGCCGCCTCCACGCAGAGCGCGCCTCTGGCCTTGGTCACTTCCCTCGCGACTACAGTATCTGGTCCCGGCC
GCGGCGGCAGCGGCGGCGGGGGGACCAGTAGCGGGGCGAGCCGTAGCTGCAGTCCCGCATCCTCGGAGGCCACTGCAGCGC
CCGGTGACCGGCTGAGAGCTGAGAGCCCGTCGCCCCCACGCTTGCTGGCTGCACACTGCGCGCTGCTGCCCAAGCCCGGATTC
CTGGGCGCCGGAGGAGGCGGCGGCGCGGCGGGTGGGCCGGGCACTCCCCACCACCACGCGCACCCTGGTGCAGCAGCCGCC
GCGGCTGCCGCTGCCGCTGCCGCGGCTGCCGGTGGCCTGGCACTGGGGCTGCACCCGGGGGGCGCACAGGGCGGCGCGGGC
CTCCCTGCACAGGCGGCTCTCTATGGACACCCGGTCTACAGTTATTCGGCAGCAGCTGCAGCGGCCGCGCTAGCTGGCCAGCA
CCCGGCGCTTTCCTACTCATACCCTCAGGTGCAGGGCGCGCACCCTGCGCACCCTGCCGACCCCATCAAGCTGGGTGCCAGCA
CCTTCCAACTGGACCAGTGGCTGCGCGCGTCTACTGCGGGCATGATCCTGCCCAAGATGCCGGACTTCAGCTGTGAGTACCTGC
GCCGCACCCGGACAGTTCTCACGCGCATCCTCCTCTCTCGCACCCCCTTCCTTTTGCTCTCACCTCCTAGCGAGCTGTGGAAGAG
CATCGAAGCCTCTGTGGCTCATTCCTGGGCCTGTGAGGCCTCTCGTGGGCCGCACAGAATGAAAGGAGCCTCAAATTATTAAGA
TGGCCACAAGCAGGCCTTGGCCTCTCTATCTCTCTTAACAGACAGGGGAGGTTGGAGGTGTGAGGCTAAAACCTAAAGAATAA
CCTGCAAGCTGGACAAGGAAGTGCGCGGAAAGGCTAGCCTCGATTAGGGCAGGTTTTGGGGGTCGCTGAAACCGGGGACTTAT
CTTGAGAAAATGAACTTTGTCAGAGGTTCCATTATAGGCTATGAAGAGACACATCACTGGGTCCAGAAGTAGCTCTTCCGTGGC
CCTTGTAAGGACTCTGCCTTCCTTTGAAGGGCAAGACATTTGCATGTCTCTGTCTCTCTGTCTGTCTCTGTCTGTCTCTGTCTCTGT
CTCTGTCTCTGTCTCTCTCTCTCTCTCTGTGTGTGTGTGTGTGTGTGTGTGTGTGTGTGTGTGTGTTGGTGAGGGTGGGCGTCCCTC
CAAATGATGTCGAACTGGTCTGTTTGAATATAAGGTCACCGGGTCTATAAATAGAAAGTGCCTGGCTGGGAACTTCAGGTGCTC
AACACCTTGGGGAGTTCCTGTCCAAGGTGAGGGCACAAAGGTCCAGTCTTCTGCATCCTCTCTGTTACCCAATAGGGACAACCT
TGCGGGAGGGGGGACATTTCCGACTTCCCTGGCACATGCAGCGTCTGTGTAGGGCACCCTGGCTCTGATTTCTGCTCTGTGCCT
GACTTGGATAGGGGCAAGCTGCCTTTCCTGAATGGATGGACCAGAAGATGTATATGGCTTTCTCCTGTCAGCCACTCAACACCC
AAACAAAGAGACCCCATTGTTTCCGTCTGAGTGGGTTTGTTTAGACTGGGAAGATGCAGGCACTGGAGGGAGAAAAAAAGCAA
AAAAGAAAACAGAAATAAACATTTGCCAGAATTCAGATTCCTCTGCTGGAAGCGGTGGTCATGACTAAACAGAGAAAGCTGGG
GCCCCAGAGAACCGCAGAAGTAGCAAGGTGGTGAGAGAATCACAGCCTGCGCACTTGTTTGCTCGCAGCCACCCAGGTTTTTC
CCAGCTTGGCTCCAAGAAGACTAGCCTTGTTGATCTCAGCAAAGAGGAAAGGCTCCCTTTAACCTCAGTTGGACTGCAGCTCAG
ACATCTCTGAAATCATTCCTATTCACTAAGTCCTAGGCAGGATCTGTGCCTGGCTGAGGAACCCAGGTTTCAGATGTCCACTATG
GCTGGACTTTCCAGGCCTGAGAAGCATAGTGGTCCTGAGAAGAGCTGACCAGTGGTCAACAACAGGCCTCCATAGCTCTGGGT
TTACTGGCTACACAAGTGGTCTCTGGGAGAAAAGGGATGTTTTTGTGGAGCCCAGGATCAATCAAAATCCATGTAAGTGGGATC
CCTTTGCCAGAATATCCAGCCCATGATGTCTCATGGTGTGGGCATTTAAAGTATGGCCAGGTCTACATTAGGAAAAGAAAGCCA
AGCCCCAGATCCCAATGGACACTAAGGAATGCAGCGGAGGGTGGCTGAATATGTTCTGTTCCTGGGACTTCCCTCTCCCCTGGG
AAAACTAGCTTGGTTTTCTTTAATTGGTTGTTTCTACTGGATGGCATTTCCCAAATGATCCCTTCCAAGACAGCGTTTTAATTAGAG
TTCTAATGTATTGTCAAGCTGGGAATTTGTAGCCTTGTTTAACTAAAGCAGGGGGCGTGGACCTTCCTATAAAGCAAATTACCCA
TCCCAGGGGAGCCTGAGCTCATCCCAGCCAGGGGCTGGCCTCTTTCTTTCCCTTGCTTTCTTTCTTTTATTGTCTAACACTTTTCGC
ACTCCTGGGAAATCAAGGGAAGTTTCATTTTGAGGTCATAAAATTCTCTGTGATGTAACTGTGCGAAATTATGTAAATAAAGCTG
GACAGATCTCCCGGAACACACAGAAAGCCAGATCTACTCACAGCTCAACCTTTAATTTGAGTAATGAGAATCGTCCCAATAGGC
TTCCAGTCCTCAGCCTTTGGCCTTAGCACTAGGAGGGGAGAGTTTGTCAACAGAGACGTGAAGACACGCAGATGAATGCCTATT
TAGACTCTAGCTTCAGTCCCTATCCTTCCTTTTCTCCCTCCCCCACCGTTCCTGCGCCCACGCCCCCTGTCTCTTCTGCTTTAAGAT
CGTGGTGGAACCAAGAGGATGGGTTCTTTTCCAGCTTGAGGTGGGAGAAAGCCTTCCATGAGAATGGGAGCGACCTGGTCTCT
ATGCTTGCTAGTCCAGTTTAGCTACCCAAGCCTTTAGACCAACCCTCCTCCTGTTCATTCGGGTGGCCACTGAGCGGAGGCAAC
AGGCCCTTTTTTCTCAAATGAGATGAGCTCGCCACTTCCACTTCATGTCTGAGATGTGGGCCATAGGGAGTGGGCACTGAGGCT
GAAGCGTCTGGAGGGTGGGCTCTGGAGGTTGGATCACGGGAGGGAAAAGGGCAGCGACGGGTGGAACACTACCTCGCCCTCA
GTGGACAAATTTACAGAAGCCTCTTGCCCTTTCTGTCCCCAGCCCAGGCGCAGTCGAACCTCTTGGGGAAGTGCCGAAGGCCTC
GCACGGCCTTCACCAGCCAGCAGCTGTTGGAGCTGGAACACCAGTTCAAGCTCAACAAGgatcctctagagtcgacctgcaggcatgcaa
gcttggcgtaatcatggtcatagctgtttcctgtgtgaaattgttatccgctcacaattccacacaacatacgagccggaagcataaagtgtaaagcctggggtgcctaatga
gtgagctaactcacattaattgcgttgcgctcactgcccgctttccagtcgggaaacctgtcgtgccagctgcattaatgaatcggccaacgcgcggggagaggcggtttg

cgtattgggcgc
